# Chance and Necessity in the Pleiotropic Consequences of Adaptation for Budding Yeast

**DOI:** 10.1101/724617

**Authors:** Elizabeth R. Jerison, Alex N. Nguyen Ba, Michael M. Desai, Sergey Kryazhimskiy

## Abstract

Mutations that a population accumulates during evolution in one (“home”) environment may cause fitness gains or losses in other conditions. Such pleiotropic fitness effects determine the evolutionary fate of the population in variable environments and can lead to ecological specialization. It is unclear how the pleiotropic outcomes of evolution are shaped by the intrinsic randomness of the evolutionary process and by the deterministic variation in selection pressures across environments. To address this question, we evolved 20 replicate populations of the yeast *Saccharomyces cerevisiae* in 11 laboratory environments and measured their fitness across multiple other conditions. We found that evolution in all home environments led to a diversity of patterns of pleiotropic fitness gains and losses, driven by multiple types of mutations. Approximately 60% percent of this variation are explained by clone’s home environment and the most common parallel genetic changes, while about 40% are attributed to the stochastic accumulation of mutations whose pleiotropic effects are unpredictable. On average, populations specialized to their home environment, but generalists also evolved in almost all conditions. Our results suggest that the mutations accumulating in a home environment incur a variety of pleiotropic effects, from costs to benefits, with different probabilities. Therefore, whether a population evolves towards a specialist or a generalist phenotype is heavily influenced by chance.

## Introduction

Populations adapt by accumulating mutations that are beneficial in their current environment, along with linked hitchhiker mutations [1]. If the population finds itself in a new environment, the effects of these accumulated mutations may change, potentially conferring fitness benefits in the new condition or incurring fitness costs. Such byproduct (or pleiotropic) effects of adaptation in one condition on fitness in others can expand the organism’s ecological niche [2–4], lead to ecological specialization and speciation [4–6] and help maintain genetic and phenotypic diversity in populations [7, 8]. Fitness trade-offs can also be exploited for practical purposes, for example, to create attenuated antiviral vaccines [9], slow down the evolution of multi-drug resistance [10], and offer the opportunity to use fluctuating drug treatments to reduce the probability of drug failure [11]. However, despite decades of research, we still lack a fundamental understanding of the statistical structure of pleiotropy, especially for new mutations [3, 6–8, 12–15]. That is, how do mutations that arise and reach high frequencies in a population adapting to one condition typically affect fitness in other conditions?

Historically, it has been assumed that pleiotropy is often antagonistic, i.e., fitness benefits in one environment should often come at a fitness cost in other conditions [16–18]. If antagonistic pleiotropy was common, it would explain why ecological specialization and local adaptation are so widespread in nature. However, more recent field studies have found that adaptive alleles confer fitness defects much less frequently than anticipated [8, 13–15, 19]. Although theory suggests that ecological specialization and local adaptation can arise without trade-offs [20–22], it is also possible that field studies provide a skewed view of the structure of pleiotropy because of statistical complications and confounding factors, such as migration and unknown environmental variation [23–25].

Laboratory microbial and viral populations are powerful model systems where the structure of pleiotropy can be probed under controlled conditions and with a degree of replication seldom achievable in natural systems. Experimental populations can be evolved in a laboratory environment, adaptive mutations can be identified, and the fitness of evolved genotypes can be directly measured in other conditions. Several dozen such studies in a variety of organisms have been carried out so far (e.g., Refs. [26–45] and more references in recent reviews by Elena [15] and Bono et al [14, 22]). Their outcomes generally support the conclusions from the field that fitness trade-offs exist [26–29, 31–40, 42, 45, 46] but are not ubiquitous [30, 32, 34, 37, 41, 43, 44, 47].

The reasons for the differences in outcomes of adaptation in different evolution experiments are not entirely clear [14, 19, 37]. One possibility is that the pleiotropic outcomes depend primarily on the differences in selection pressure between the home and the non-home environments [37, 42]. In other words, it is possible that adaptation to any given home environment leads to the accumulation of mutations with some typical, home-environment dependent, fitness effects in other conditions. Then, the observed differences in pleiotropic outcomes of evolution would be primarily determined by the choice of home and non-home environments. It is also possible that chance events play an important role [14, 34, 37]. Different mutations may have dramatically different pleiotropic effects [14]. Since independently evolving populations acquire different sets of mutations, we would expect to observe different pleiotropic outcomes even among populations that evolved in the same condition. In this case, the probability that any particular population evolves towards a specialist or a generalist phenotype would depend both on the environmental selection pressures and on the distribution of pleiotropic effects of available beneficial mutations.

Disentangling and quantifying the contributions of selection and chance to the pleiotropic outcomes of adaptive evolution is difficult even in the laboratory because it requires observing evolution in many replicate populations and measuring their fitness in many other conditions. To this end, we evolved populations of yeast *Sacchromyces cerevisiae* in a variety of laboratory environments, sequenced their full genomes, and measured the fitness of the evolved clones in multiple panels of non-home conditions. Due to the intrinsic randomness of evolution, populations evolving in the same condition accumulate different mutations. Thus, to quantify the contribution of natural selection and evolutionary stochasticity to pleiotropy, we estimate the variance in the pleiotropic fitness gains and losses explained by these two factors. In addition, we examine how pleiotropic outcomes depend on the similarity between the new and the home environments.

## Results

To investigate the pleiotropic consequences of adaptation, we experimentally evolved 20 replicate *S. cerevisiae* populations in 11 different laboratory environments (a total of 220 populations). Each population was founded from a single colony isolated from a common clonal stock of a laboratory strain. We chose the 11 laboratory environments to represent various degrees of several different types of physiological stresses (e.g. osmotic stress, temperature stress). A complete list of all 11 evolution conditions, plus two additional conditions used for assays, is provided in Table 1.

**Table 1.** Environmental conditions used in this study.

| Environment | Evolution Condition | Measurement Panels |  |  |  | Formulation |
| --- | --- | --- | --- | --- | --- | --- |
|  |  | Diagnostic | Salt | pH | Temp |  |
| SC | ✓ | ✓ | ✓ | ✓ | ✓ | SC + 2% glu, 30°C |
| Low Salt | ✓ |  | ✓ |  |  | SC + 2% glu + 0.2 M NaCl, 30°C |
| Med Salt |  |  | ✓ |  |  | SC + 2% glu + 0.4 M NaCl, 30°C |
| High Salt | ✓ | ✓ | ✓ |  |  | SC + 2% glu + 0.8 M NaCl, 30°C |
| pH 3 | ✓ | ✓ |  | ✓ |  | SC + 2% glu, buffered to pH 3, 30°C |
| pH 3.8 | ✓ |  |  | ✓ |  | SC + 2% glu, buffered to pH 3.8, 30°C |
| pH 6 | ✓ |  |  | ✓ |  | SC + 2% glu, buffered to pH 6, 30°C |
| pH 7.3 | ✓ | ✓ |  | ✓ |  | SC + 2% glu, buffered to pH 7.3, 30°C |
| Low Temp | ✓ | ✓ |  |  | ✓ | SC + 2% glu, 21°C |
| Med Temp |  |  |  |  | ✓ | SC + 2% glu, 34°C |
| High Temp | ✓ | ✓ |  |  | ✓ | SC + 2% glu, 37°C |
| Low Glu | ✓ | ✓ |  |  |  | SC + 0.07% glu, 30°C |
| Gal | ✓ | ✓ |  |  |  | SC + 2% gal, 30°C |

SC is synthetic complete medium; glu is glucose; gal is galactose.

We evolved each population in batch culture at an effective size of about *N_e_ ≈* 2 × 10^5^ for about 700 generations using our standard methods for laboratory evolution (see Methods for details). Seven populations were lost due to pipetting errors during evolution, leaving a total of 213 evolved lines. We randomly selected a single clone from each evolved population for further analysis.

### Specialization is the typical outcome of adaptation

To understand how adaptation to one (“home”) environment alters the fitness of the organism in other (“non-home”) environments, we measured the competitive fitness of each evolved clone relative to their common ancestor across multiple conditions (Methods). We first focused on a “diagnostic” panel of eight conditions that represent different types of physiological stresses (see Table 1).

Figure 1 shows the median change in fitness of these clones across the eight diagnostic conditions. As expected, clones evolved in all environments typically gained fitness in their home environment, although the magnitude of these gains varied across conditions (diagonal entries in Figure 1). We quantified the degree of specialization of clones evolved in a given home environment as the average fraction of non-home environments where clones lost fitness relative to their ancestor (Methods). Figure 1 (left bar graph) shows that, by this definition, populations evolved in all environments typically specialized, but the degree of specialization varied between home environments. For example, evolution at pH 3 caused a typical population to lose fitness in all but one non-home diagnostic environment, whereas evolution at Low Temp did not cause a typical population to suffer significant fitness losses in any of the diagnostic environments.

**Fig 1.**
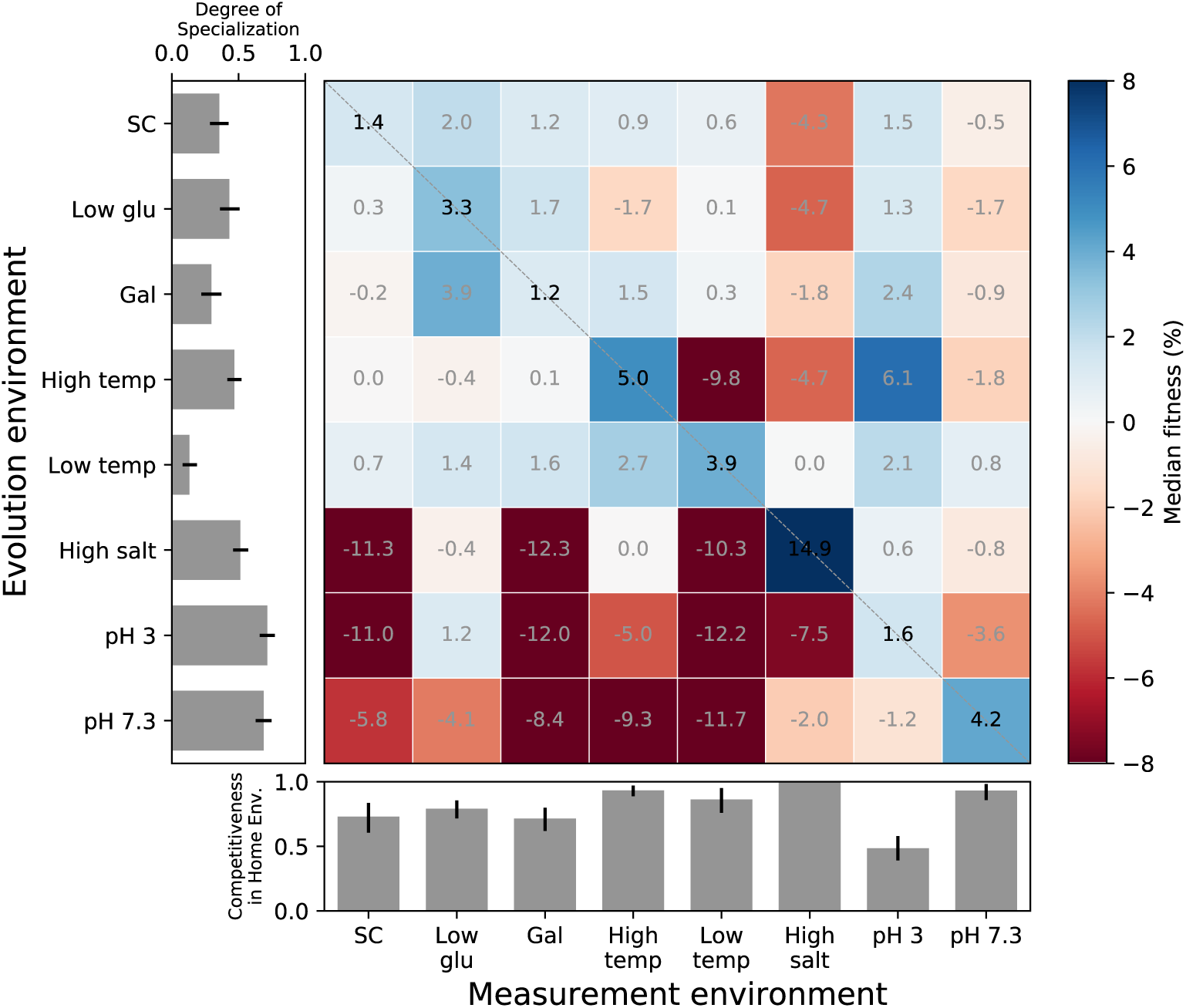
Fitness gains and losses in diagnostic conditions after evolution in each condition. Each square shows the median fitness gain or loss in a measurement environment (columns) across all populations evolved in a given home environment (rows) for ∼700 generations. The left bar graph shows the average degree of specialization after evolution in each home environment. The degree of specialization is measured as the average proportion of measurement environments where clones lost fitness relative to the ancestor. The bar graph on the bottom shows the competitiveness of a “resident” clone in its home environment against invasions from other environments. The competitiveness is measured as the average proportion of evolved clones from other environments that are less fit than a randomly chosen resident clone. Error bars represent 95% confidence intervals on a bootstrap over clones in each evolution condition.

For the long-term survival of an ecological specialist, its fitness in the home environment must be higher than that of populations evolved elsewhere. To test whether adaptation to a given environment leads a “resident” population to become a better competitor in its home environment than “invader” populations evolved elsewhere, we estimated the proportion of pairwise competitions between residents and invaders where the resident wins (Methods). We found that, in most home environments, an average resident is able to outcompete most or all invaders from other environments (Figure 1, bottom bar graph). The exception to this rule is the pH 3 environment, where residents lost in more than half of competitions.

We conclude that adaptive evolution typically leads populations to specialize to their home environment, and the evolved specialists are typically able to resist invasions from populations evolved elsewhere. As expected, the specific set of conditions where an evolved population gains and loses fitness depends on the population’s home environment. One exception is the unexpected similarity between pleiotropic consequences of evolution in three apparently unrelated conditions: adaptation to High Salt, pH 3 and pH 7.3 led to similar and large median fitness losses in SC, Gal, and Low Temp.

### Evolution leads to diverse but environment-specific pleiotropic outcomes

The patterns of median pleiotropic fitness gains and losses shown in Figure 1 may be driven by differences in selection pressure between environments, such that mutations acquired in different environments have systematically different pleiotropic effects in other conditions. Alternatively, these patterns could have arisen because each clone stochastically acquired a different set of mutations and each set of mutations produces its own idiosyncratic pattern of pleiotropic fitness gains and losses across environments.

To discriminate between these two possibilities, we quantified the variation in the patterns of pleiotropic fitness gains and losses around the medians observed in Figure 1. For each clone, we calculated its “pleiotropic profile”, the 8-dimensional vector containing its fitness changes (relative to the ancestor) in the eight diagnostic environments. If clones isolated from the same home environment cluster together in this 8-dimensional space, it would indicate that evolution in this environment leaves a stereotypical pleiotropic signature. Lack of clustering would suggest that the patterns in the median pleiotropic profiles shown in Figure 1 are driven by evolutionary stochasticity and idiosyncratic pleiotropy.

To visualize the clustering of pleiotropic profiles, we used t-stochastic nearest neighbor embedding (t-SNE) to project the 8-dimensional profiles onto two dimensions (Figure 2A,B). This t-SNE embedding is useful in looking for cluster structure because it minimally distorts the original local distances between points, such that clones that are close together in the two-dimensional embedding have similar 8-dimensional pleiotropic profiles (in contrast to principal components analysis where this may not always be the case). In Figure 2D, we show the patterns of pleiotropy associated with each of the measured clones. Each clone is colored consistent with its color in Figure 2B.

**Fig 2.**
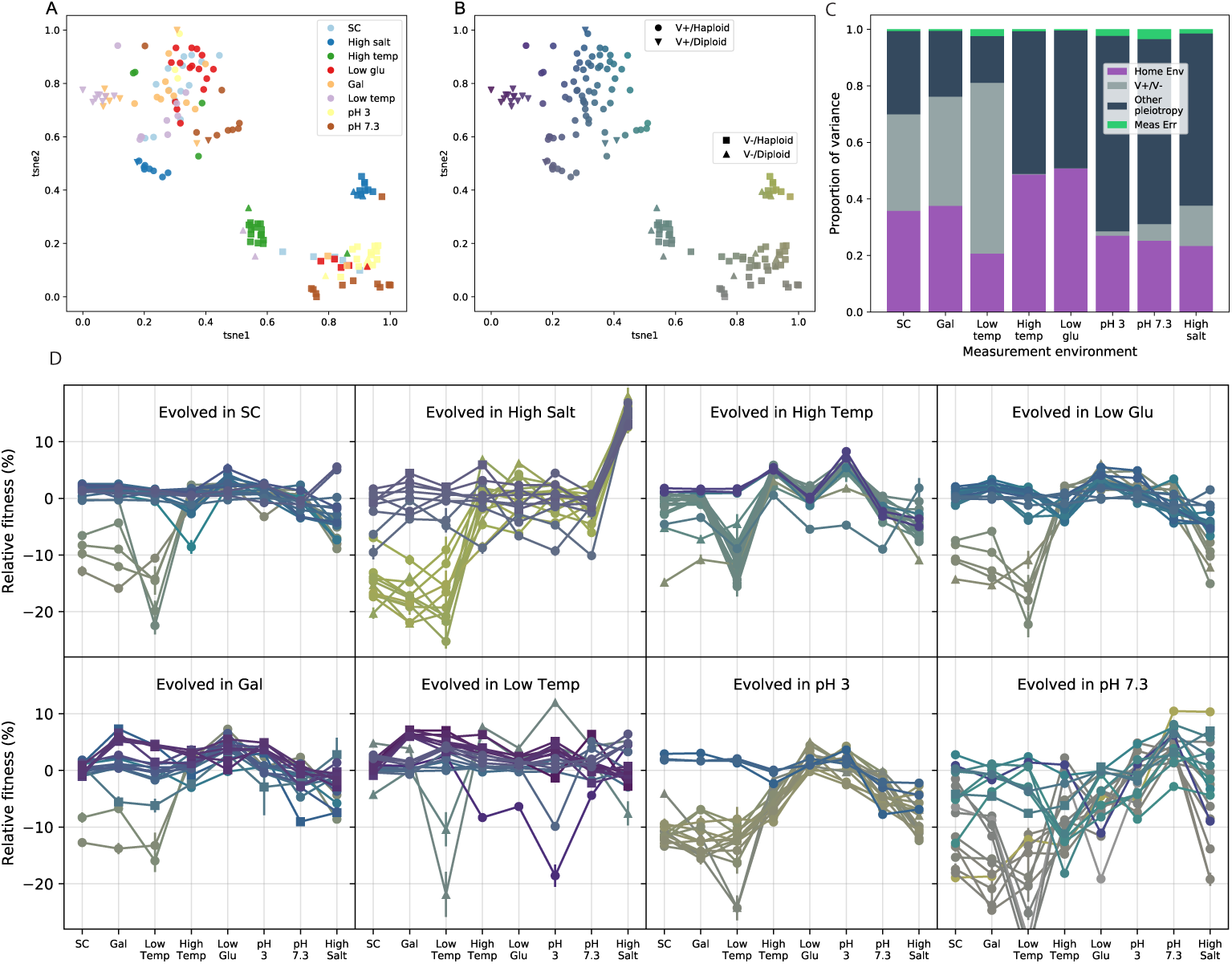
Environmental and stochastic determinants of pleiotropic profiles. **A.** t-SNE dimensional reduction of the pleiotropic profiles. Each point represents a clone; the eight-dimensional vector of clone fitness across the eight conditions was projected into two dimensions using t-SNE. Clones are colored according to home environment. **B.** t-SNE projection as in A. Colors are assigned based on the t-SNE coordinates to establish visual correspondence between this projection and the full pleiotropic profiles shown in panel D. **C.** Proportion of the variance in clone fitness in each environment that is attributable to (in this order) home environment, V^+^/V*^−^*phenotype, other pleiotropy (unexplained variance), measurement noise. Clones are excluded from their own home environment. **D.** The pleiotropic profiles of clones from each home environment. Profiles are colored as in B. Error bars represent *±*1 SE on clone fitness.

The t-SNE embedding reveals that there are two large and clearly separated clusters, both of which contain clones from all home environments (Figure 2A). The main features that discriminate the two clusters are the fitness in SC, Gal and Low Temp (Figure 2B,D). Clones that belong to one cluster lost 10 to 40 percent in these conditions, whereas clones that belong to the other cluster did not (Figure 2B,D). We refer to these two phenotypes as V*^−^* and V^+^, respectively, for reasons that will become clear in the next section.

Beyond these two large clusters, the distribution of clones evolved in different conditions in the t-SNE space is not uniform. First, clones from some home environments are more likely to have the V*^−^*phenotype than clones from other environments (*χ*^2^-test, *p* = 6.8 ×10*^−^*^8^). For example, 16/20 clones evolved in High Temp are V*^−^* but only 3/20 clones evolved in Gal are V*^−^*. In fact, this non-uniform distribution of V^+^ and V*^−^* clones among home environments explains the large median fitness losses in the SC, Gal and Low Temp conditions exhibited by clones evolved in High Salt, pH 3 and pH 7.3 which we saw in Figure 1 (see also Figure S1). Second, although there is a substantial overlap between the distributions of clones evolved in different home environments within the large V^+^ and V*^−^* clusters (Figure 2A), clones from some environments clearly form tight smaller clusters (e.g., High Salt clones). In general, neighbors of a clone are more likely to be from the same home environment: on average, 2.8 of the 5 nearest neighbors are from the same environment, compared with 0.60 ± 0.12 under random permutation.

Next, we set out to quantify the extent to which the observed variation in pleiotropic profiles is explained by the deterministic differences in selection pressures between environments versus the intrinsic randomness of the evolutionary process. Using a nested linear model, we estimated the fractions of observed variance in fitness in each diagnostic environment that is attributed to the identity of the home environment of a clone and to measurement noise. We attribute the remaining, unexplained, variance to evolutionary stochasticity, i.e., the fact that each clone acquired a unique set of mutations which have idiosyncratic pleiotropic effects. We found that the home environment accounts for between 20% and 51% of the variance in fitness, depending on the diagnostic environment (Figure 2C). Measurement noise accounts for less than 4% of variance, leaving 48% to 77% unexplained, i.e., attributable to evolutionary stochasticity (Figure 2C). However, if, in addition to clone’s home environment, its status with respect to the V^+^/V*^−^*phenotype becomes known (for example, after measuring its fitness in another condition, such as Low Temp), the fraction of unexplained variance drops to 16%–70%, depending on the diagnostic environment (Figure 2C).

Taken together, these observations show that the home environment leaves a distinct signature in the clone’s pleiotropic profile, such that clones evolved in the same condition tend to be more similar to each other than clones evolved in different conditions. However, these deterministic differences are generally less important than the randomness of the evolutionary process, accounting for on average 34% of the variance in pleiotropic outcomes, compared with 65% for stochastic effects.

### The genetic basis of pleiotropic outcomes

Next, we sought to determine the genetic basis underlying the diverse pleiotropic outcomes that we observed above, using two approaches. First, we used DNA staining and flow cytometry (see Methods) to look for ploidy changes because this is a common mode of adaptation in yeast [48–52]. Second, we sequenced the full genomes of the evolved clones. We carried out these analyses on all 213 clones, i.e., those evolved in the diagnostic conditions considered above as well as other intermediate-stress environments listed in Table 1. Sequencing failed at the library preparation stage or due to insufficient coverage in 15 cases, leaving us with 198 sequenced clones. Using standard bioinformatic methods for calling SNPs and small indels (Methods), we identified a total of 1925 *de novo* mutations. We note that, because our sequencing and analysis pipeline can result in false negatives (i.e. certain mutations are difficult to confidently identify), our results represent a subset of all mutations in each sequenced clone.

### Loss of killer virus causes the V***^−^*** phenotype

We began by looking for the genetic differences between the V^+^ and V*^−^* clones. We found no association between V^+^ or V*^−^* phenotypes and ploidy or any of the mutations identified in the sequencing data. Instead, multiple lines of evidence demonstrate that the V*^−^*phenotype was caused by the loss of the yeast killer virus, a toxin-antitoxin system encoded by a ~ 2 kb cytoplasmic dsRNA [53–57] that was present in the ancestor of our experiment (and was retained in the V^+^ clones).

First, we directly looked for the presence or absence of the corresponding band in a gel electrophoresis assay (Methods). We found that both the ancestor and 7 out of 7 randomly selected V^+^ clones displayed the killer-virus band, while all of the 7 randomly selected V*^−^* clones did not (Figure 3A). Second, we cured the ancestor strain of the killer virus (Methods) and competed all our evolved clones against this new cured reference strain at Low Temp, which we chose as a test environment because the fitness defect is largest in this condition. We observed that the severe fitness defect that the V*^−^* clones have at Low Temp when they compete against their direct ancestor entirely disappears in competitions against the cured ancestor (Figure 3B). In addition to these two experiments, we obtained several other pieces of evidence (see Methods and Figures S2–S4) which support the conclusion that the loss of the killer virus is the cause of the V*^−^* phenotype.

**Fig 3.**
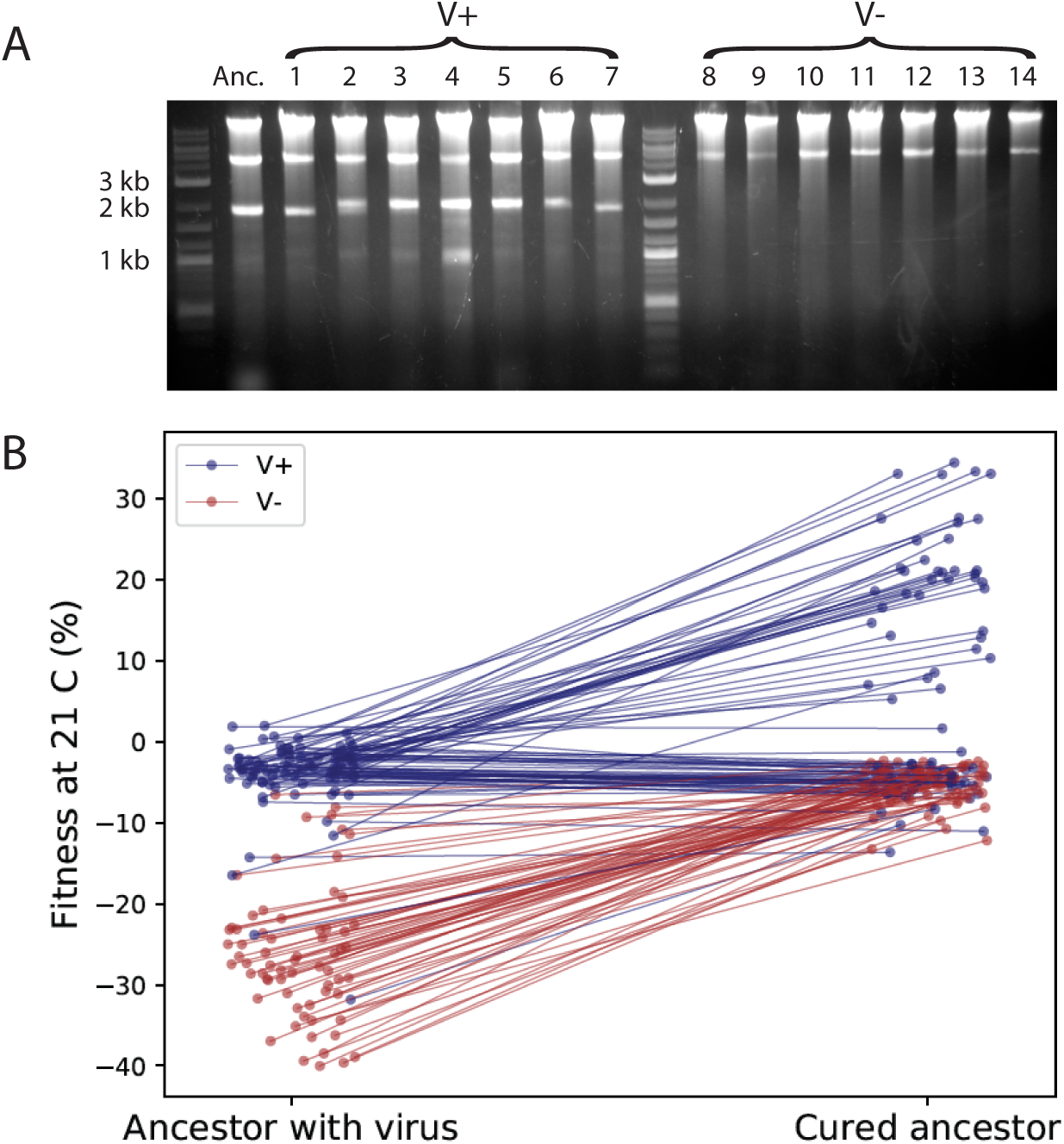
V*^−^* phenotype is caused by the loss of yeast killer virus. **A.** Gel electrophoresis of total DNA and dsRNA extracted from 15 clones. Anc is the common ancestor of the experiment; evolved clones 1 through 7 are from the V^+^ cluster (Figure 2), i.e., without a fitness defect at 21°C; evolved clones 8 through 14 are from the V*^−^* cluster, i.e., with a fitness defect at 21°C. The upper (∼ 4 kb) band is consistent with the helper virus, and the lower (∼ 2 kb) band is consistent with the killer virus. **B.** Fitness of all evolved clones relative to the ancestor and to the ancestor cured of the killer virus. Classification of clones into V^+^(blue) and V*^−^*(red) is based on the t-SNE plot in Figure 2B.

Our results suggest that the severe fitness defects in SC, Low Temp, and Gal environments (Figures 1 and 2D) are not due to an inherent growth disadvantage. Rather, V*^−^* clones suffer large losses of fitness in competitions against the virus-carrying ancestor because they succumb to the virus expressed by the ancestor. Consequently, these fitness losses are frequency-dependent (Figure S4). They are particularly severe in SC, Low Temp, and Gal likely because virus activity is higher in these conditions [58]. Nevertheless, virus loss evolved even in these environments (Figure 2D). This initially puzzling observation could be explained if virus virulence was lost first and resistance was lost second, after non-virulent genotypes dominated the population. In support of this explanation, we found some evolved clones that have similar fitness relative to both the virus carrying and virus-cured ancestors (Figure 3B, horizontal lines), suggesting they are resistant but non-virulent [59]. A recent study that examined the co-evolution of yeast and its killer virus also reported such stepwise progression towards virus loss and showed that virus loss likely provides no fitness benefit to the host [60].

### Diversity at the genetic level underlies diversity of pleiotropic outcomes

We next looked for the genetic basis for the fine-scale phenotypic variation between clones that we observed in our t-SNE plot (Figure 2A,B). We found that 35 out of 213 clones became diploid during evolution. Diploids evolved more often in some environments than in others (*p* = 1.3 *×* 10*^−^*^4^, *χ*^2^-test) and 24 out of 35 diploids retained the killer virus, while 11 lost it (Figure 2B). Moreover, 13 V^+^ diploid clones that evolved in Low Temp and Gal formed a small cluster in the t-SNE space (Figure 2A, inverted triangles), suggesting that a change in ploidy, irrespective where it evolved, leads to certain characteristic changes in the pleiotropic profile, perhaps in conjunction with other mutations.

We next used our full-genome sequencing data to call putatively beneficial SNPs and indels. We identified such mutations as nonsynonymous, nonsense, or frameshift changes within “multi-hit” genes, defined here as genes that were mutated in four or more clones across all home environments or in two or more clones from the same home environment (see Methods and Figure S5). In total, we identified 176 such mutations in 42 multi-hit genes (Figure 4A). Only three individual multi-hit genes (SIR3, HNM1, and PDE2) were significantly associated with one home environment (*p <* 0.01, Bonferroni-corrected permutation test; see Methods). Mutations in most other multi-hit genes arose in multiple home environments, but with significantly different frequencies (*p <* 10*^−^*^4^, Methods; Figure 4A).

**Fig 4.**
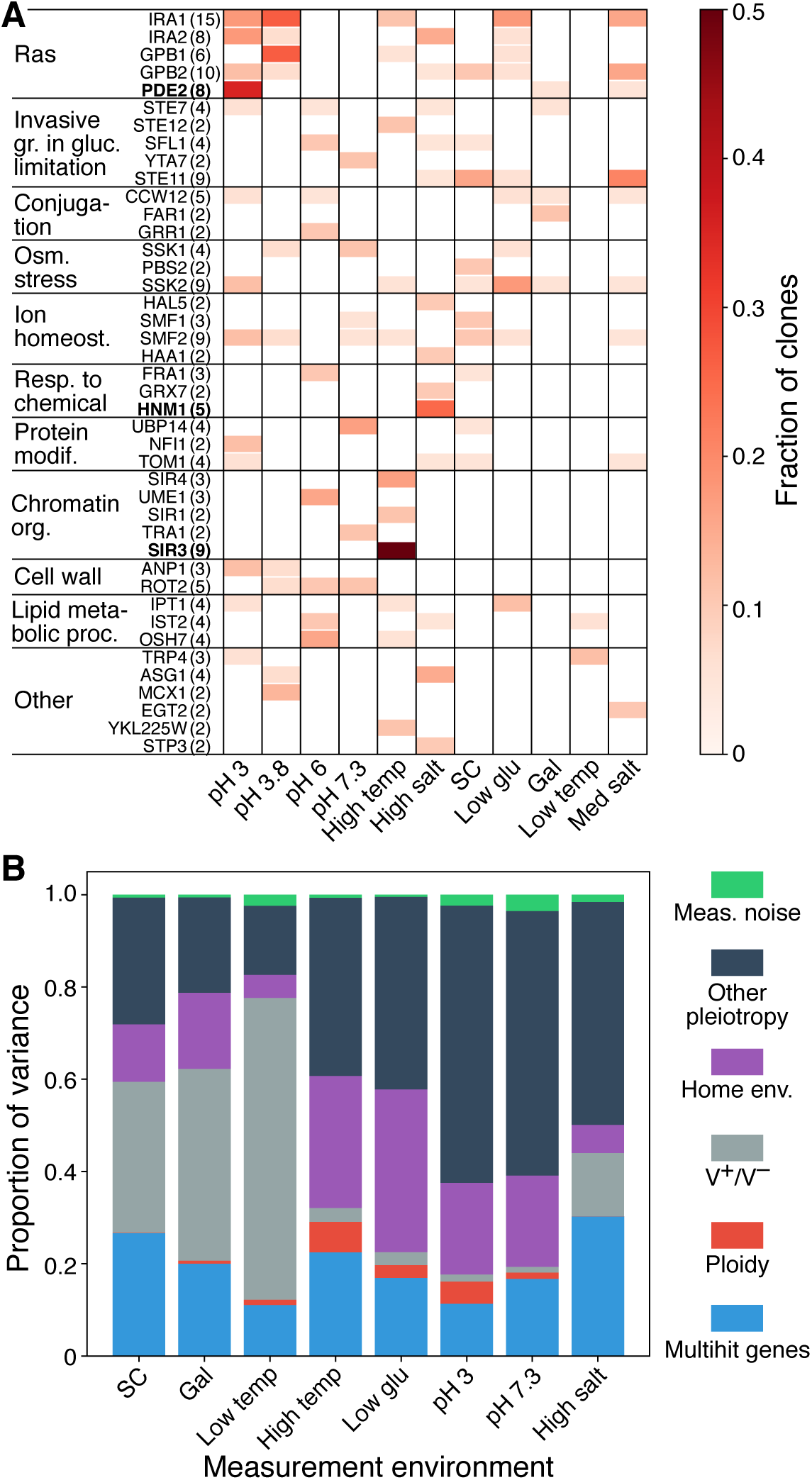
Mutations across evolution conditions and genetic determinants of pleiotropy. **A.** Genes with four or more nonsynonymous mutations across the experiment, or two or more within one home environment, organized into the Ras pathway and thereafter by GO slim process. Number in the parentheses next to each gene name indicates the total number of detected nonsynonymous mutations in that gene. Genes in bold are significantly associated with a single home environment **B.** Proportion of the variance in clone fitness in each environment attributable (in this order) to mutations in multi-hit genes; ploidy; V^+^/V*^−^* phenotype; home environment, beyond these previously listed factors; other pleiotropy (unexplained variance); measurement noise. Clones are excluded from their own home environment. Only clones evolved in diagnostic conditions are considered in this analysis, as in Figure 2.

To quantify the extent to which this genetic information improves our ability to statistically predict the fitness of a clone in a diagnostic environment, we expanded the list of predictor variables in the nested linear model described in the previous section to include the presence or absence of multi-hit mutations shown in Figure 4A and the ploidy status. We found that mutations in multi-hit genes account for 11%–30% of the variance in pleiotropic effects (Figure 4B), and all genetic factors combined account for 17%–77% of variance. After accounting for these genetic factors, clone’s home environment still explains 5%–35% of variance. This implies that, even though mutations in some genes fail to meet the multi-hit gene significance threshold, they nevertheless have somewhat predictable pleiotropic effects. After accounting for all these factors, the fraction of unexplained variance drops to 15%–60%, which we now attribute to the accumulation of passenger mutations with unpredictable pleiotropic effects.

In summary, multiple types of genetic changes accumulate during evolution in all our environments, including point mutations in a diverse set of target genes, diploidization and killer-virus loss. The same genetic changes often occur in populations evolving in different environments, which leads to substantial uncertainty in the outcomes of evolution at the genetic level. Nevertheless, the probabilities of observing any given type of mutation are different in different environments, such that the genotype of a clone at a relatively few loci explains about half of variation in the pleiotropic outcomes of adaptation. Together with the knowledge of population’s home environment, genetic differences on average explain approximately 60% percent of this variation. The remaining ∼ 40% are attributed to the stochastic accumulation of mutations whose pleiotropic effects are unpredictable.

### Fitness trade-offs are not inevitable but their frequency increases with dissimilarity between environments

We observed that clones adapted to a home environment concomitantly gain fitness in some non-home conditions and lose fitness in some other non-home conditions (Figures 2 and S1). We also established that these patterns of pleiotropic gains and losses depend on the home environment. We next sought to understand what determines whether a clone evolved in one condition gains or loses fitness in another. Our hypothesis is that fitness is pleiotropically gained in conditions that are in some sense similar to the home environment and lost in conditions that are dissimilar to the home environment [37, 43]. Testing this hypothesis in our original diagnostic panel of 8 environments is difficult because it is not clear how similar or dissimilar they are. Therefore, we focused on three panels of environments where in each panel yeast is exposed to a particular type of physiological stress: salt, temperature or pH stress (see Table 1). Within each panel, the test environments differ by the intensity of that stress, so that similarity between conditions within a panel is well defined.

In this analysis, we included clones evolved in all of our evolution conditions, including some intermediate-stress home environments that were not part of the diagnostic panel (see Table 1). To simplify interpretation, we restricted this analysis to V^+^. To include only V^+^ clones from intermediate-stress home environments, we measured their fitness relative to the original and cured reference strains at Low Temp, and used behavior in this assay as a classifier (Methods, Figure S6).

We first asked whether the physico-chemical similarity between environments explains the average patterns of pleiotropic fitness gains and losses. Analogously to Figures 1 and S1, we found that clones usually gained more fitness on average in their home environment than clones evolved in other conditions in the same panel (Figure 5A–C). The mean fitness of a clone steadily declined in conditions progressively less similar to its home environment. These results support the hypothesis that the similarity between environments determines the patterns of pleiotropy, at least on average. Higher moments of the distribution of pleiotropic outcomes might also depend on the similarity between conditions, but the patterns are less clear (Figure S7).

**Fig 5.**
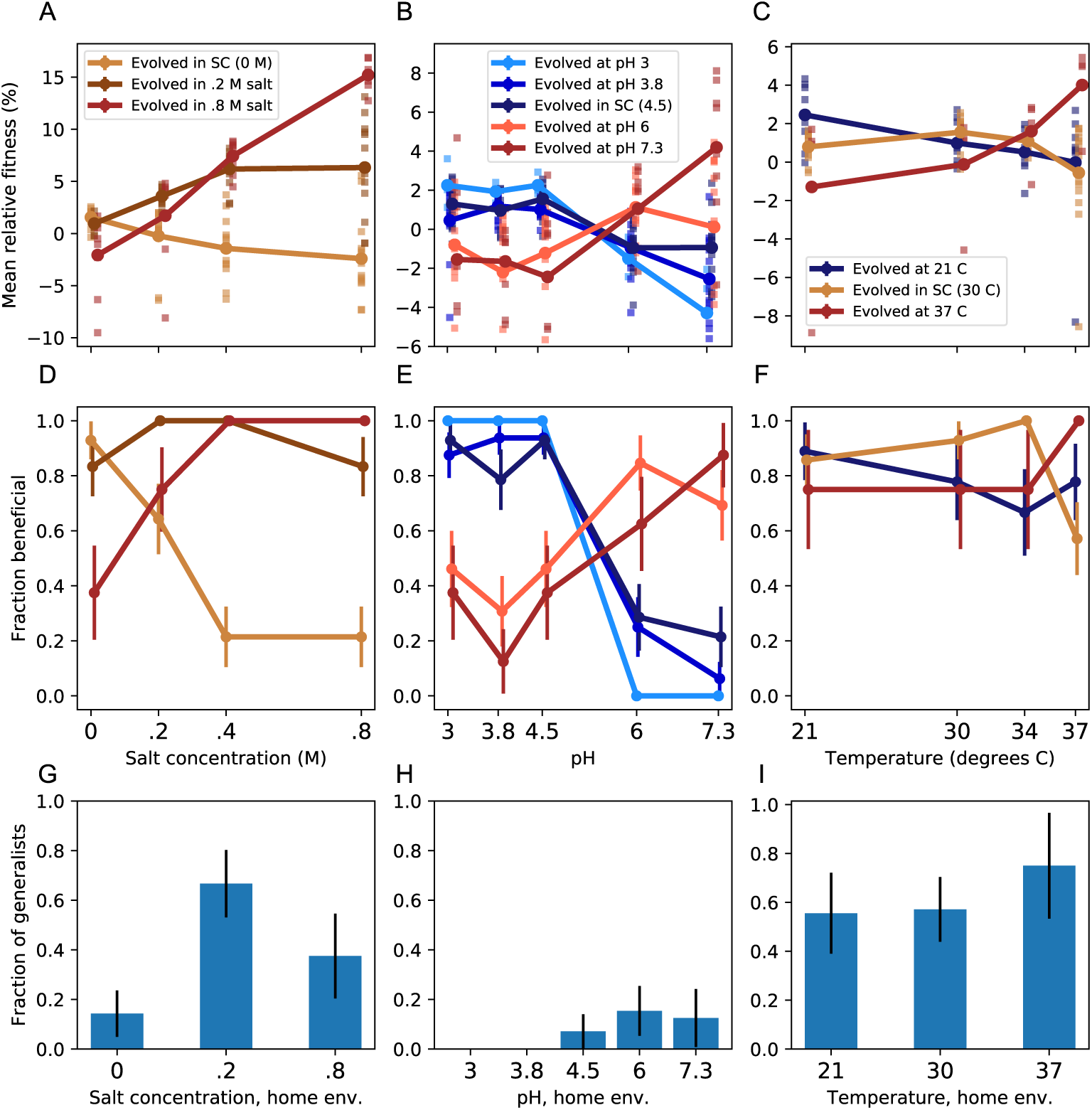
Specialization across salt, pH, and temperature panels of environments. Panels A, D, G refer to the salt panel; panels B, E, H refer to the pH panel; and panels C, F, I refer to the temperature panel. **A-C.** Average fitness of clones from each home environment (colors) across multiple test conditions (*x*-axis). Squares represent individual clone fitness. **D-F.** Fraction of clones from each home environment (colors) that gained fitness in each test condition (*x*-axis). Error bars represent ±1 SE. **G-I.** Fraction of clones from each home environment (*x*-axis) that gained fitness across all test conditions in the panel. Error bars represent *±*1 SE.

The fact that clones evolved at one extreme of a panel on average lost fitness at the other extreme (Figure 5A–C) suggests that there may be inherent physiological trade-offs between fitness in dissimilar environments. However, we found that many clones evolved at one extreme of each panel actually gained fitness at the other extreme of the panel (Figure 5D–F). For example, while the majority of V^+^ clones evolved in SC experienced fitness declines in High Salt, 3/15 experienced fitness increases, including two clones with fitness improvements of about 5% (Figure 5A). The only exception were the clones evolved in the more acidic environments all of which lost fitness in the most basic conditions (Figure 5B,E). However, some of the clones evolved in the more basic environments gained fitness in the more acidic conditions. In fact, we found a number of clones from a variety of home environments within each panel that improved in fitness across the entire panel (Figure 5G–I). While such generalist clones arise in almost all environments, clones that gain fitness in less similar environments are less common than clones that gain fitness in more similar conditions (Figure 5D–F).

These results demonstrate that there exist mutations that are beneficial across the entire range of environments that vary along one physico-chemical axis. Thus, the trade-offs between fitness even in the most dissimilar conditions (along one environmental parameter axis) are not physiologically inevitable. To further corroborate this conclusion, we measured the correlation between fitness of clones in pairs of environments in each panel (Figures S8–S10). If fitness trade-offs between a pair of conditions were physiologically inevitable, we would expect a negative correlation between fitness measured in these conditions. Instead we observe diverse and complex fitness covariation patterns, but there is a notable lack of strong negative correlations between clone fitness even in the most dissimilar pairs of environments. In conclusion, our results suggest that whether a population evolves towards a specialist or a generalist phenotype depends on the specific set of mutation that it accumulates, i.e., this outcome is largely stochastic.

## Discussion

To assess how chance and necessity in evolution affect the fitness of an organism across multiple environments, we evolved populations of budding yeast in a variety of laboratory “home” conditions. We characterized each population by its “pleiotropic profile”, the vector of fitness gains and losses in an array of diagnostic environments. We found that a diverse set of pleiotropic profiles arose during evolution in all home conditions. Underlying this phenotypic diversity, we found a diversity of evolutionary outcomes at the genetic level. Despite this large diversity, the home environment leaves statistically distinct signatures in the genome, which, in turn, cause the clones from the same home environment to have statistically similar pleiotropic profiles. We estimated that clone’s home environment and the set of most common genetic changes together explain about 60% of variance in clone’s pleiotropic fitness gains and losses. The remaining ∼ 40% are attributable to evolutionary stochasticity, i.e., the accumulation of hitchhikers or rare beneficial variants whose pleiotropic effects are unpredictable.

Despite the fact that the pleiotropic outcomes of evolution in any individual population are to a large degree governed by chance, clear and repeatable patterns emerge when we consider ensembles of populations evolved in the same home environment. For example, on average, evolution leads to specialization, so that individual’s pleiotropic fitness gains are smaller or turn into losses in environments that are more dissimilar from the home environment (Figure 5). The most obvious explanation for these repeatable patterns is that different environments exert different selection pressures on the organism. However, we cannot exclude the possibility that there may also be different spectra and rates of mutations in different environments, as a recent report shows can occur in yeast [61]. We conclude that subtle differences in selection pressures and possibly mutational biases shift the statistical patterns of pleiotropy and the distribution of fixed mutations, leading to an average degree of specialization that depends on the dissimilarity between environments.

Our results help us better understand the evolution of specialists and generalists, a long-standing problem in evolutionary ecology [12, 22, 62]. To explain the ubiquity of specialists, many models require physiological trade-offs, or antagonistic pleiotropy, so that mutations that increase fitness in one environment necessarily come at a cost of reducing fitness in other environments [16, 19, 62]. On the other hand, it has long been appreciated that fitness losses in non-home environments can arise without physiological trade-offs if the population accumulates mutations that are neutral in the home environment and deleterious elsewhere [20, 43]. However, field and experimental studies so far failed to find a pattern clearly favoring one model over another [3, 8, 13, 14].

To explain existing data, Bono et al recently proposed a model that unifies the antagonistic pleiotropy and mutation accumulation perspectives [14]. They suggest that the fitness effects of mutations in a particular pair of home and non-home environments form a continuum, such that some of the mutations that accumulate in the home environment (both drivers and hitchhikers) incur pleiotropic fitness costs in the non-home conditions and some others provide pleiotropic fitness benefits (see Figure 1 in Ref. [14]). If mutations that incur pleiotropic costs are more common and are more beneficial in the home environment than those that provide pleiotropic benefits, the populations will tend to lose fitness in the non-home condition.

Our results are consistent with the general model proposed by Bono et al. They also suggest that the pleiotropic consequences of adaptation are highly stochastic, i.e., they depend on the particular set of mutations that the population happened to have accumulated during evolution in the home environment. Moreover, the chance for a population to acquire a pleiotropically costly or pleiotropically beneficial mutation is not the same in all environments. If the non-home environment is physico-chemically similar to the home environment, the fitness effects of mutations in the two conditions will be strongly correlated. As the similarity declines, more mutations that are beneficial in one become deleterious in the other. As a result, populations are more likely to suffer pleiotropic fitness costs in conditions more dissimilar from the home environment.

In this work, we examined the statistics of pleiotropy among beneficial mutations that arise in populations of a particular size descended from one particular ancestral yeast genotype. These statistics likely depend on the population size because populations of different size sample different sets of adaptive mutations [63]. It is also likely that different genotypes have access to beneficial mutations with different statistics of pleiotropy [64]. To understand these broader patterns, we need to know the joint distribution of fitness effects of new mutations and how it varies across genotypes due to epistasis.

Assuming that the structure of pleiotropy does not change drastically between closely related genotypes, our results would suggest that longer periods of evolution in a constant environment would lead to further specialization, simply because pleiotropically costly mutations are more abundant. If the environment fluctuates between two or multiple states, adaptive mutations that are less common but provide fitness gains in multiple conditions would spread giving rise to generalist genotypes. Why then do “jacks of all traits” not evolve? Our results suggest that such genotypes may not be physiologically impossible but are simply extremely unlikely because mutations that are beneficial in larger and larger sets of distinct conditions are too rare for populations with realistic sizes to discover them.

## Materials and methods

### Experimental evolution

The *S. cerevisiae* strain yGIL104 (derived from W303, genotype *Mat**a**, URA*3*, leu*2*, trp*1*, CAN*1*, ade*2*, his*3*, bar*1Δ *:: ADE*2, [65]) was used to found 220 populations for evolution. Each population was founded from a single colony picked from an agar plate. Populations were propagated in batch culture in 96-well polystyrene plates (Corning, VWR catalog #29445-154), with 128*µl* of media per well. Populations evolving in the same environment were grown in wells B2-B11 and E2-E11 on the same plate. Except for the galactose and low glucose conditions, all media contained 2% dextrose (BD, VWR catalog #90000-904), 0.67% YNB with nitrogen (Sunrise Science, catalog #1501-500), and 0.2% SC (Sunrise Science, catalog #1300-030). The galactose condition contained 2% galactose (Sigma-Aldrich, #G0625) instead of dextrose, and the low glucose condition contained 0.07% dextrose. Other conditions contained the following in addition to SC-complete. Low salt: 0.2 M sodium chloride. Medium salt: 0.4 M sodium chloride. High salt: 0.8 M sodium chloride. pH 3: 0.02 M disodium phosphate, 0.04 M citric acid. pH 3.8: 0.0354 M disodium phosphate, 0.032 M citric acid. pH 6: 0.0642 M disodium phosphate, 0.0179 M citric acid. pH 7.3: 0.0936 M disodium phosphate, 0.00032 M citric acid. Buffered media were filter sterilized; all other media were autoclaved.

All populations were grown at 30°C, except for the high temperature lines (37°C) and the low temperature lines, which were grown at room temperature (21 ± 0.5°C). In the SC, high temperature, medium salt, low glucose, pH 3, pH 3.8, and pH 6 conditions, dilutions were carried out once every 24 hours. In the galactose, low temperature, and high salt conditions, dilutions were carried out every 36 hours. All dilutions were carried out on a Biomek-FX pipetting robot (Beckman-Coulter). Before each transfer, cells were resuspended by shaking on a Titramax 100 orbital plate shaker at 1200 rpm for at least 1 minute. In the pH 7.3 condition, dilutions were carried out every 48 hours. At each transfer, all populations were diluted 1:512 except for the low glucose populations, which were diluted 1:64. This maintained a bottleneck size of about 10^4^ cells in all conditions. Populations underwent approximately the following numbers of generations (doublings): SC, high temperature, medium salt: 820. Low glucose: 730. pH 3, pH 3.8, pH 6: 755. High salt, galactose, low temperature: 612. Every 7 transfers, populations were mixed with glycerol to final concentration 25% (w/v) and stored at −80°C. Each 96-well plate contained blank wells; no contamination of blank wells was observed during the evolution. Over the course of evolution, 7 populations were lost due to pipetting errors, leaving 213 evolved lines.

To pick clones for further analysis, each final population was streaked onto SC-complete with 2% agar. One colony per population was picked, grown in 128 *µl* of SC at 30°C, mixed with 25% (w/v) glycerol, and stored at *−*80°C.

### Competitive fitness assays

We conducted flow-cytometry-based competitive fitness assays against yGIL104-cit, a fluorescently-labeled derivative of the common ancestor, yGIL104. To construct the fluorescent reference strain, we amplified the HIS3MX6-ymCitrineM233I construct from genomic DNA of strain yJHK111 (courtesy of Melanie Muller, John Koschwanez, and Andrew Murray, Department of Molecular and cellular Biology, Harvard University) using primers oGW137 and oGW138 and integrating it into the his3 locus. The fitness effect of the fluorescent marker is less than 1% in all environments (Figure S11).

Fitness assays were conducted as has been described previously [66, 67]. Briefly, we grew all test strains and the reference strain from frozen stock in SC at 30°C. After 24 hours, we diluted all lines into the assay environment for one growth cycle of preconditioning. We then mixed the reference strain and the test strains 50/50. We monitored the relative numbers of the reference and test strain over three days in co-culture. We measured fitness as 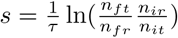 where *τ* is the number of generations between timepoints, *n_it_* is the count of the test strain at the initial timepoint, *n_ft_* is the count of the test strain at the final timepoint, and *n_fr_*, *n_ir_* are the counts for the reference.

### Library preparation and whole-genome sequencing

Libraries were prepared for sequencing as has been described previously [68]. Briefly, genomic DNA was extracted from each of the 213 clones using the PureLink Pro 96 Genomic Purication Kit (Life Technologies catalog #K1821-04A) and quantified using the Qubit platform. The multiplexed sequencing library for the Illumina platform was prepared using the Nextera kit (Illumina catalog #FC-121-1031 and #FC-121-1012) and a modified version of the Illumina-recommended protocol [68]. Libraries were sequenced on a Nextera Hi-seq 2500 in rapid-run mode with paired-end, 150 bp reads.

### Nucleic acid staining for ploidy

Clones were grown to saturation in YPD (2% dextrose, 2% peptone, 1% yeast extract). Saturated cultures were diluted 1:10 into 120 *µl* of sterile water in a 96-well plate. The plate was centrifuged, and cultures were resuspended in 50 *µl* of fresh water. 100 *µl* of ethanol was added to each well, and the wells were mixed slowly. Plates were incubated for one hour at room temperature, or overnight at 4°C. Cells were centrifuged, ethanol solution removed, and 65 *µl* RNAase solution added (2 mg/mL RNAase in 10 mM Tris-HCl, pH 8.0, plus 15 mM NaCl). Samples were incubated at 37°C for 2 hours. To stain, 65 *µl* of 300 nM SYTOX green (ThermoFisher, S-34860) in 10 mM Tris-HCl was added to each well, for a final volume of 130 *µl*. Plates were incubated at room temperature, in foil, for 20 minutes.

Fluorescence was measured via flow cytometry on a Fortessa analyzer (FITC channel). Fluorescence peaks were compared to known haploid and diploid controls to score ploidy. For 19/213 clones we observed smeared peaks intermediate between the haploid and diploid peaks; we called these clones as undetermined and included them with the haploids in analysis.

### SNP and indel identification

We called SNPs and small indels as described previously [69], with the following two modifications: first, we aligned reads to a custom W303 reference genome [70]. Second, for clones called as diploid via staining, we called mutations as heterozygous if they occurred at frequencies between 0.4 and 0.8, and homozygous otherwise. We called mutations in all other clones if they occurred at a frequency of at least 0.8. We included both heterozygous and homozygous mutations in subsequent analyses.

For 95.4% (1925) of the mutations that we called, the mutation was found in one clone (i.e. the mutation was unique at the nucleotide level). The remaining 4.6% (88) of mutations were found in two or more clones. These mutations may have originated from standing genetic variation in the starting strain, and thus we excluded them from our analysis of *de novo* mutations.

### Analysis of genetic parallelism

To test for parallelism at the gene level, we redistributed the observed nonsynonymous mutations across the genes in the yeast genome, under a multinomial distribution with probabilities proportional to the gene lengths. We determined that genes with four nonsynonymous mutations across the experiment, or two nonsynonymous mutations within one evolution condition, were enriched (Figure S5). To divide these genes into categories, we first classified genes as belonging to the Ras pathway based on *de novo* mutations in the same pathway found in previous studies [50, 70]. We classified the remainder of the genes using GO-SLIM ‘biological process’ analysis, placing genes into GO-SLIM categories in order of the process enrichment score.

To test for associations between individual multi-hit genes and home environments, we redistributed the observed mutations in each gene across environments, preserving the number of mutations per gene and the number of mutations per environment, but ignoring which mutations occurred in which clones. We calculated the nominal P-value by comparing the maximum number of hits to a particular gene in any environment in the permuted and original data. To correct for multiple testing, we multiplied the obtained nominal P-value by the total number of genes (Bonferroni correction).

We used a mutual-information based test statistic to test for overall association between the evolution environments and mutated genes. We defined the mutual information as:

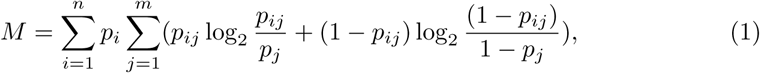

where *m* is the number of significant genes, *n* is the number of evolution environments, *p_ij_* is the probability of a clone from environment *i* having a mutation in gene *j*, *p_j_* is the probability of any clone having a mutation in gene *j*, and *p_i_* is the proportion of clones evolved in environment *i*. By convention *p_ij_* log_2_(*p_ij_*) = 0 if *p_ij_* = 0, and probabilities were estimated based on the observed frequencies of the events. We determined significance by comparing *M* to the null distribution under permutation, preserving the number of mutations per gene and the number of mutations per environment. For the null distribution, *M* was .67 (.62, .73), whereas for the data *M* was 1.15. Code used for analysis and figure generation is available at: https://github.com/erjerison/pleiotropy. The number of sequenced clones from each environment was: SC (19), High Salt (20), High Temp (18), Low Glu (17), Gal (18), Low Temp (18), pH 3 (17), pH 7.3 (18), pH 6 (19), pH 3.8 (15), Med Salt (19).

### t-SNE and clustering analysis

We used the sklearn.manifold.t-SNE class in the Python package scikit-learn 0.2, with 2 dimensions and perplexity 30, to project the 8-dimensional fitness vectors into a dimensional t-SNE space. We then used the sklearn.cluster.KMeans class to perform k-means clustering with k=2 in the t-SNE space. We used this cluster assignment to call V^+^ and V*^−^* phenotypes. These clusters correspond to those identifiable visually in Figure 2. The number of clones from each ‘extreme’ environment was: SC (19), High Salt (20), High Temp (20), Low Glu (20), Gal (19), Low Temp (19), pH 3 (18), pH 7.3

(20).

### Specialization and competitiveness summary statistics

To assess the degree of specialization of a clone, we counted the number of non-home environments where its fitness relative to the ancestor was 2 SEM below zero. Figure 1 (left bar chart) shows the proportion of such conditions averaged over all clones from the same home environment. To assess the competitiveness of “resident” clones in their home environment relative to clones evolved elsewhere, we estimated the proportion of all clones evolved in other conditions with fitness lower than a randomly-chosen resident clone (Figure 1, bottom bar chart). For both statistics, we measured 95% confidence intervals based on a bootstrap over clones in each evolution environment.

### Nested linear models for analysis of variance

To evaluate the fraction of the variance in pleiotropic effects attributable to the evolution condition vs. stochastic evolutionary effects, we fit the following series of nested linear models for each of the diagnostic measurement conditions:

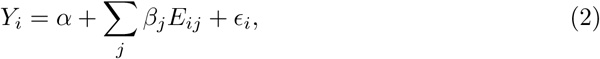

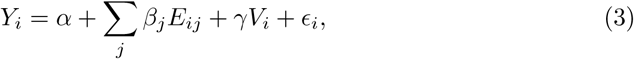

where *Y_i_* is the fitness of clone *i*; *E_ij_* = 1 if clone *i* evolved in environment *j*, 0 otherwise; *V_i_*= 1 if clone *i* is V^+^, 0 otherwise. Note that we excluded clones from their own home environment to focus on pleiotropic effects, as opposed to adaptation to the home condition. Note also that we restricted analysis to clones measured in all eight diagnostic conditions to maintain comparability between environments. We fit the models using the sklearn.linear model.LinearRegression class in Python, and used the score method of this class to calculate *R*^2^.

Figure 2C shows the partitioning of the total variance in *Y_i_* as follows. We measured the variance due to measurement error as:

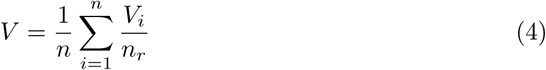

where *n* is the number of clones, *n_r_*is the number of replicate measurements of each clone, and *V_i_*is the estimate of the variance across replicate fitness measurements of clone *i*. We attribute the variance explained by model 2 to “home environment”. We attribute the variance not explained by model 2 but explained by model 3 to V^+^/V*^−^*phenotype. We attributed leftover variance not accounted for by model 3 and not attributed to measurement noise to additional stochastic effects, which we label ‘other pleiotropy.’

To evaluate the contributions of most common genetic factors to the pleiotropic effects (shown in Figure 4B), we carried out an analogous analysis of variance using the following series of nested linear models:

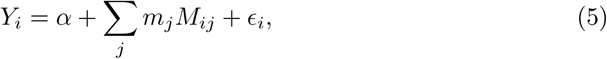

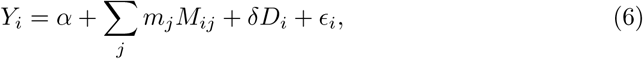

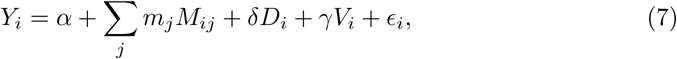

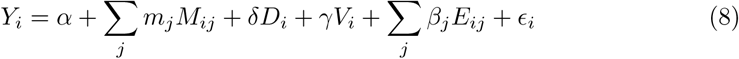

where *M_ij_* = 1 if clone *i* has at least one mutation in biological process *j* out of 10 biological processes shown in Figure 4A (other that “Other”); *D_i_* = 1 if clone *i* is a diploid, 0 otherwise.

### dsRNA extraction and gel electrophoresis

Yeast cell pellets from 1.5 mL of an overnight culture were re-suspended in 50*µ*l of a zymolyase based enzymatic digest to lyse the cells (5mg/mL Zymolyase 20T, 100mM sodium phosphate buffer pH 7.4, 10mM EDTA, 1M Sorbitol, 20mM DTT, 200*µ*g/mL RNAse A, 0.5% 3-(N,N-Dimethylmyristylammonio)propanesulfonate and incubated at 37°C for an hour. The spheroplasted cells were then lysed with 200*µ*l of lysis/binding buffer (4.125M guanidine thiocyanate, 25% isopropanol, 100mM MES pH 5). After a brief vortexing, the clear solution was passed through a standard silica column for DNA purification, and washed two times with 600*µ* l of wash buffer (20mM Tris-Cl pH 7.4, 80% ethanol). After drying the column, the DNA and dsRNA was eluted with a low salt elution buffer (10mM Tris-Cl pH 8.5).

Total extracted genomic material was subjected to standard gel electrophoresis (1% agarose gel, TAE running buffer, stained with 0.5*µ* g/mL ethidium bromide).

### Curing the killer virus

Strain yGIL104-cit-V- was constructed from yGIL104-cit as follows. yGIL104-cit was grown from frozen stock overnight in YPD. Saturated culture was diluted 1 : 10^5^ and 250 *µ* L was plated on YPD. Plates were incubated at 39°C for 72 hours. Colonies were picked and the presence of the virus dsRNA band was tested as described above. 2/9 colonies tested displayed the helper virus band but no killer virus band; 7/9 retained both bands. The two V*^−^*colonies were restreaked and a clone from each was grown in YPD, mixed with glycerol to 25%, and stored at 80°C. Competitive fitness assays were performed with both clones against yGIL104, at several starting frequencies, in SC, 21°C, 37°C, and high salt. Fitness of the two clones at each frequency and condition were the same, so one clone was designated yGIL104-cit-V- and was used as a cured reference in all subsequent assays.

We used fitness relative to the original and cured ancestor to classify clones from the 3 environments not included in the diagnostic panel as either V^+^ or V*^−^* (Figure S6). We also note that one clone (High Temp clone 20) was lost from the cured reference fitness assay.

### Determining the cause of the Low Temp fitness defect: additional experiments

We performed several types of experiments to determine the genetic basis of the large observed fitness defects in the Low Temp environment. First, we reconstructed all six nonsynonymous mutations called in one evolved clone with the fitness defect on the ancestral background. The strain background used for reconstructions was yERJ3, which was constructed from yGIL104 by amplifying the HIS3 construct from yGIL104-cit using primers 3 and 4, which target the URA3 locus. This construct was transformed into yGIL104 using standard techniques [71], plated on CSM-His dropout media, and replica plated to 5FoA and CSM-Ura to verify the His+/Ura-phenotype.

We used the delitto-perfetto method for the reconstructions [72]. Briefly, we amplified a URA3-Hph construct from plasmid pMJM37 (provided by Michael J. McDonald) using primers 6-17, which target the yeast genome 5 bp upstream and downstream of the mutations of interest. We selected on CSM-Ura and hygromycin-B, picked two clones, and transformed each with two complementary 90 bp repair oligos (18–29), that contain the mutation of interest and the flanking genic region. We selected on 5FoA and replica plated to hygromycin to determine the phenotype. We used primers 30-41 to amplify the locus in the reconstructed line for Sanger sequencing.

We performed fitness assays of yERJ3, the reconstructed lines, and the knockout intermediates, against yGIL104-cit in the SC, 37°C, 21°C, and high salt conditions. For one mutation, in the gene CUE4, one reconstruction replicate displayed a significant fitness defect across all conditions, while the other replicate did not. We discarded this clone as a likely reconstruction artifact.

We note that the reconstruction background, yERJ3, had an apparent fitness defect of a few percent in the high salt environment, potentially due to the engineered URA3 auxotrophy. We report fitness of reconstructed lines relative to yERJ3 in Figure S2. These mutations account for the fitness advantage in the clone’s home environment (High Salt), but none of them carries the characteristic large fitness defect at Low Temp.

To determine whether the defect was caused by a mutation that we did not detect during sequencing, we back-crossed three evolved clones that displayed the defect to the common ancestor and picked four-spore complete tetrads. The strain yERJ10 (genotype *Matα* yGIL104 ura3::HIS3) was constructed from yGIL104 as described above for yERJ3. The mating type was switched using an inducible Gal::HO plasmid, pAN216a-URA3-GAL::HO-Ste2pr::SkHIS3-Ste3pr::LEU2. The strain was transformed with the plasmid and plated on CSM-Ura dropout media. A colony was grown in SC-Ura dropout media with 2% sucrose overnight. 1 mL of culture was centrifuged and resuspended in SG-Ura dropout media (2% galactose) to induce. Cells were plated on SC-Leu dropout media directly after transfer to SG-Ura and 60 minutes later colonies were streaked on SD-Complete + 5FoA to eliminate the plasmid. *Matα* versions of evolved lines were constructed in the same way. After mating, diploids were selected on CSM-Ura-His dropout media. Diploids were sporulated in Spo++ media [71] plus 0.5% dextrose at 21°C for 3-5 days. Tetrads were dissected according to standard yeast genetics methods [71]. Four-spore complete tetrads from each mating were grown in SC, mixed with glycerol to final concentration 25%, and frozen at −80°C. Fitness assays of four-spore complete tetrads from each mating, competed against yGIL104-cit, were conducted as described above at 21°C. We also constructed a mitochondrial-cured version of the reference and of the evolved lines; the fitness of spores from crosses involving these lines were not distinguishable from the corresponding *ρ*+ crosses, so spore fitness were pooled.

In Figure S3, we show data from a representative one of these crosses: yERJ10 *Matα* x High Salt-17 *Mat**a*** (backcross) and High Salt-17 *Matα* x High Salt-17 *Mat**a*** (control cross). We observed that the fitness defect did not segregate 2:2, as would be expected for a Mendelian trait; rather, very few of the segregants from the back-cross displayed the defect. This observation is consistent with a cytoplasmic genetic element (the virus) that is carried by one parent (the ancestor) but not the other (evolved line), and is usually re-inherited by segregants upon mating.

Given that the defect did not appear to be caused by a nuclear genetic mutation, we next addressed whether there was evidence of a direct interaction between strains during competition. To do so, we asked whether the size of the fitness defect depended on the frequency of the competitors. In Figure S4, we show an example of such a competition experiment, between the putative virus-carrying reference and the cured ancestor at Low Temp. The strong frequency-dependence of the fitness defect is consistent with secretion of a toxin by one competitor: the strain lacking the virus (and thus the antitoxin) is at a larger disadvantage when the virus-carrying competitor is at high frequency.

Together with the direct observation of the virus band through gel electrophoresis and the competition of all of the evolved lines against the cured ancestor, as described in the main text, these observations support the conclusion that loss of the killer virus particle in some evolved lines caused the large fitness defect at Low Temp.

### Data availabilty

Data used in Figures 1,2,5 is provided in Supplementary Table S1. Data used in Figure 3 is provided in Supplementary Table S2. Data used in Figure 4 is provided in Supplementary Table S3. Code used for analysis and figure generation is available at https://github.com/erjerison/pleiotropy. The sequences reported in this paper have been deposited in the BioProject database (accession number PRJNA554163). All strains are available from the corresponding authors upon request.

## Acknowledgments

We thank members of the Desai and Kryazhimskiy labs for experimental assistance and comments on the manuscript. We thank three anonymous reviewers for thoughtful comments. MMD acknowledges support from the Simons Foundation (Grant 376196), grant DEB-1655960 from the NSF, and grant GM104239 from the NIH. SK acknowledges support from the BWF Career Award at Scientific Interface (Grant 1010719.01), the Alfred P. Sloan Foundation (Grant FG-2017-9227), and the Hellman Foundation.

## Supplementary Tables

**Supplemantary Table S1. Fitness measurement data.**

**Supplemantary Table S2. Fitness measurement data in 21°C with respect to the cured reference.**

**Supplemantary Table S3. Identified mutations.**

**Supplemantary Table S4. Primers used in this study.**

## Supplementary Figures

**Fig S1.**
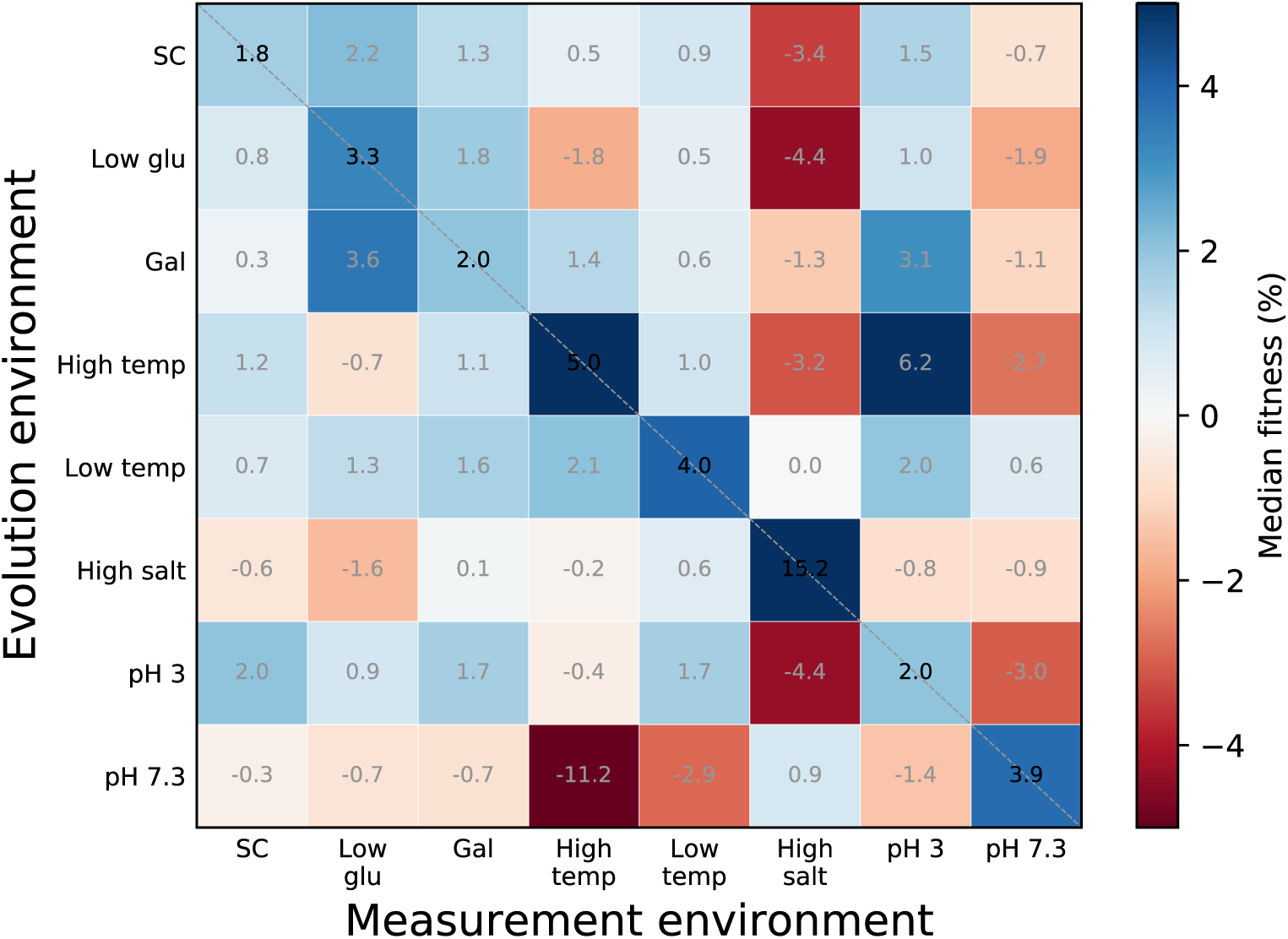
Median fitness gains and losses among groups of clones from the same home environment, excluding V*^−^* clones. Notations as in Figure 1.

**Fig S2.**
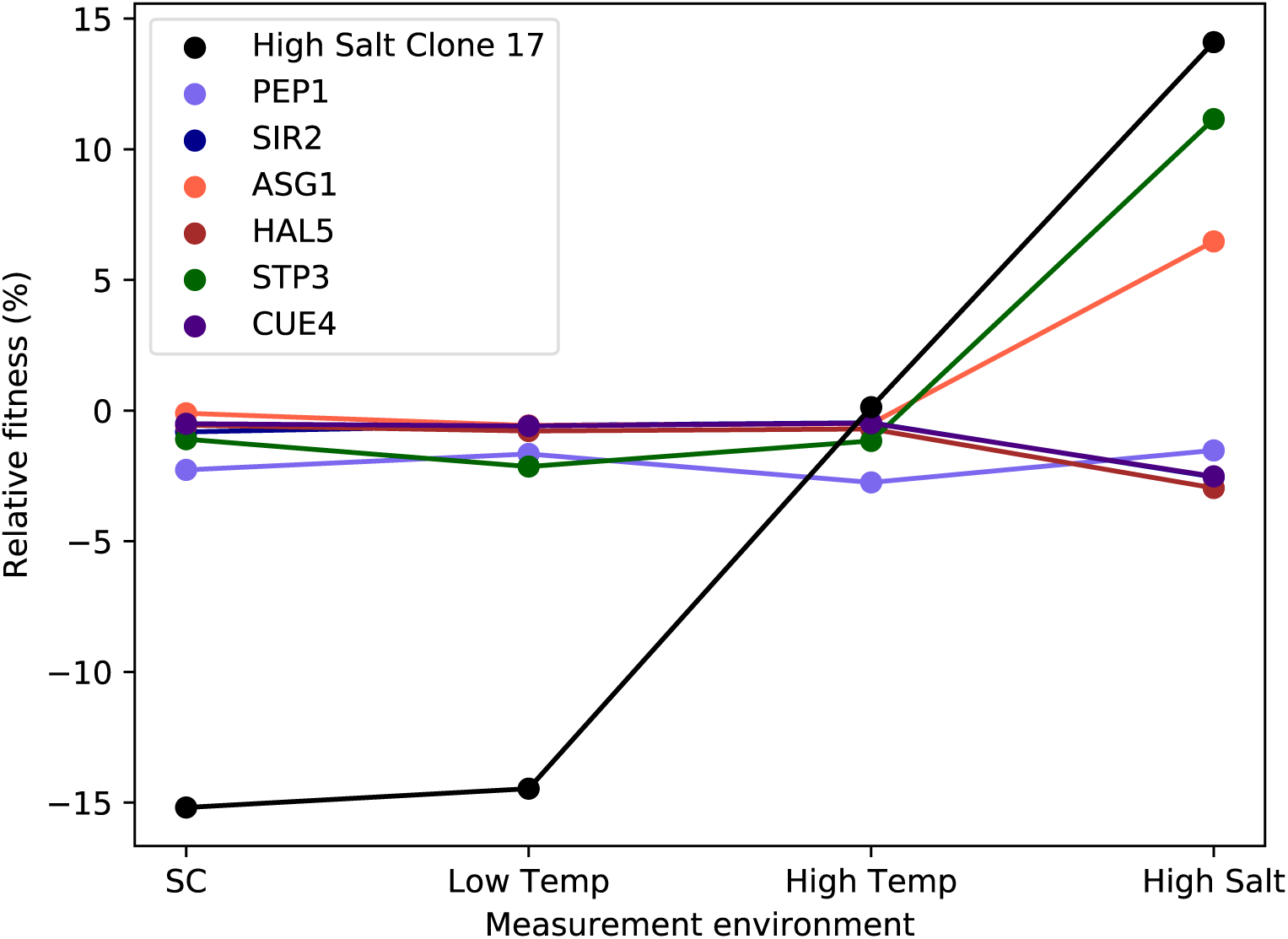
Individual reconstructions of the six nonsynonymous mutations called in one High-Salt evolved clone on the ancestral background. Fitness of the evolved clone and the six reconstructions was measured relative to the ancestor in four environments. (Error bars: *±*1 SE on reconstructed clone fitness.)

**Fig S3.**
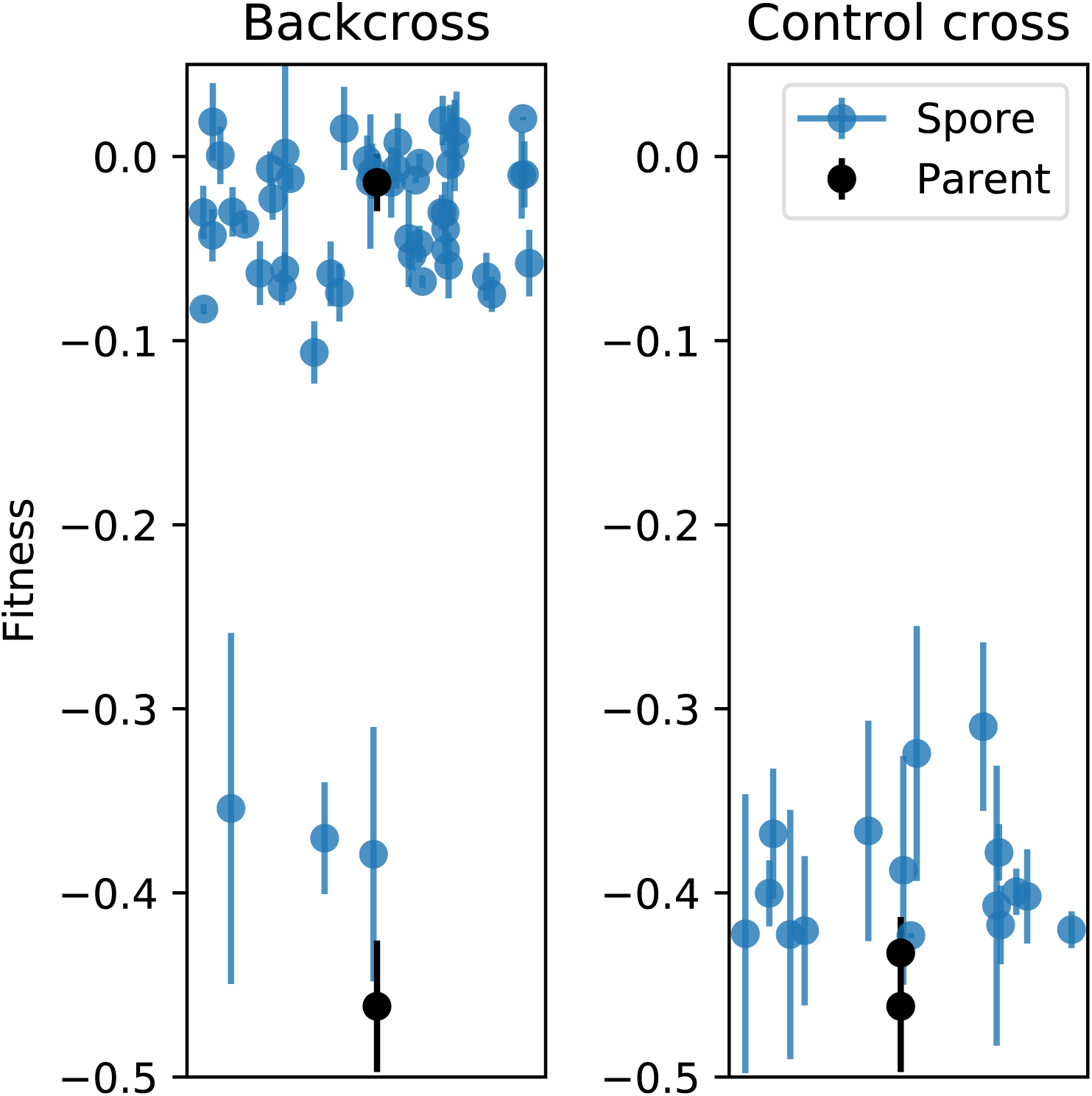
A backcross of a clone evolved in High Salt with the large fitness defect at Low Temp, to the common ancestor. Each circle represents the fitness of a spore from a four-spore complete tetrad, while squares show the fitness of the parents. (Error bars: 1 ± SE on clone fitness.) The control panel shows the cross between the evolved clone and a *Matα* version of the same genotype. The small number of clones inheriting the defect in the backcross indicates that the effect is non-Mendelian.

**Fig S4.**
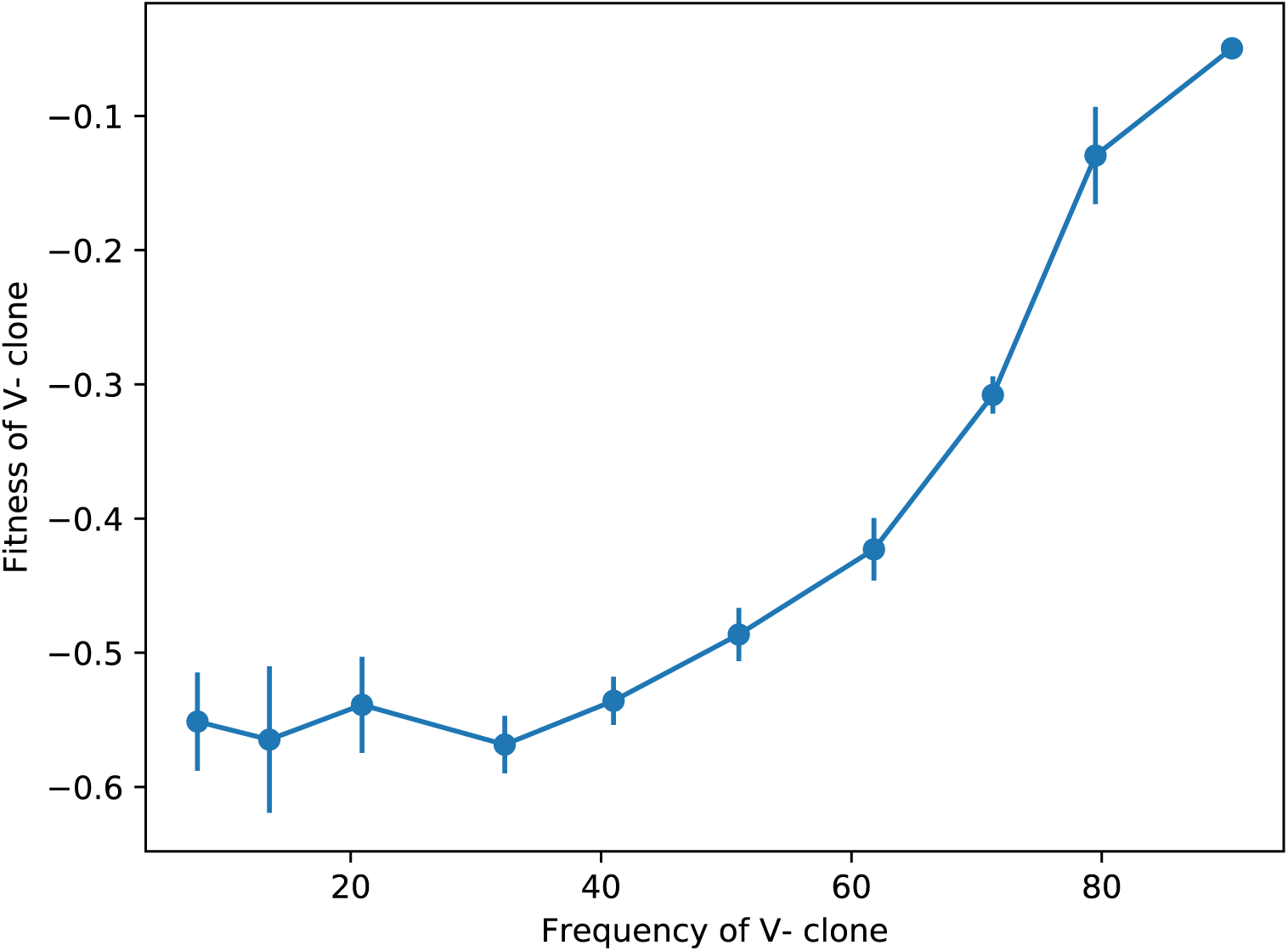
Fitness of a V*^−^* clone relative to the ancestor at Low Temp, initiated at different initial frequencies. The frequency dependence of the relative fitness suggests that the fitness defect might be caused by a direct interaction between the competitors. Error bars show *±*1 SE.

**Fig S5.**
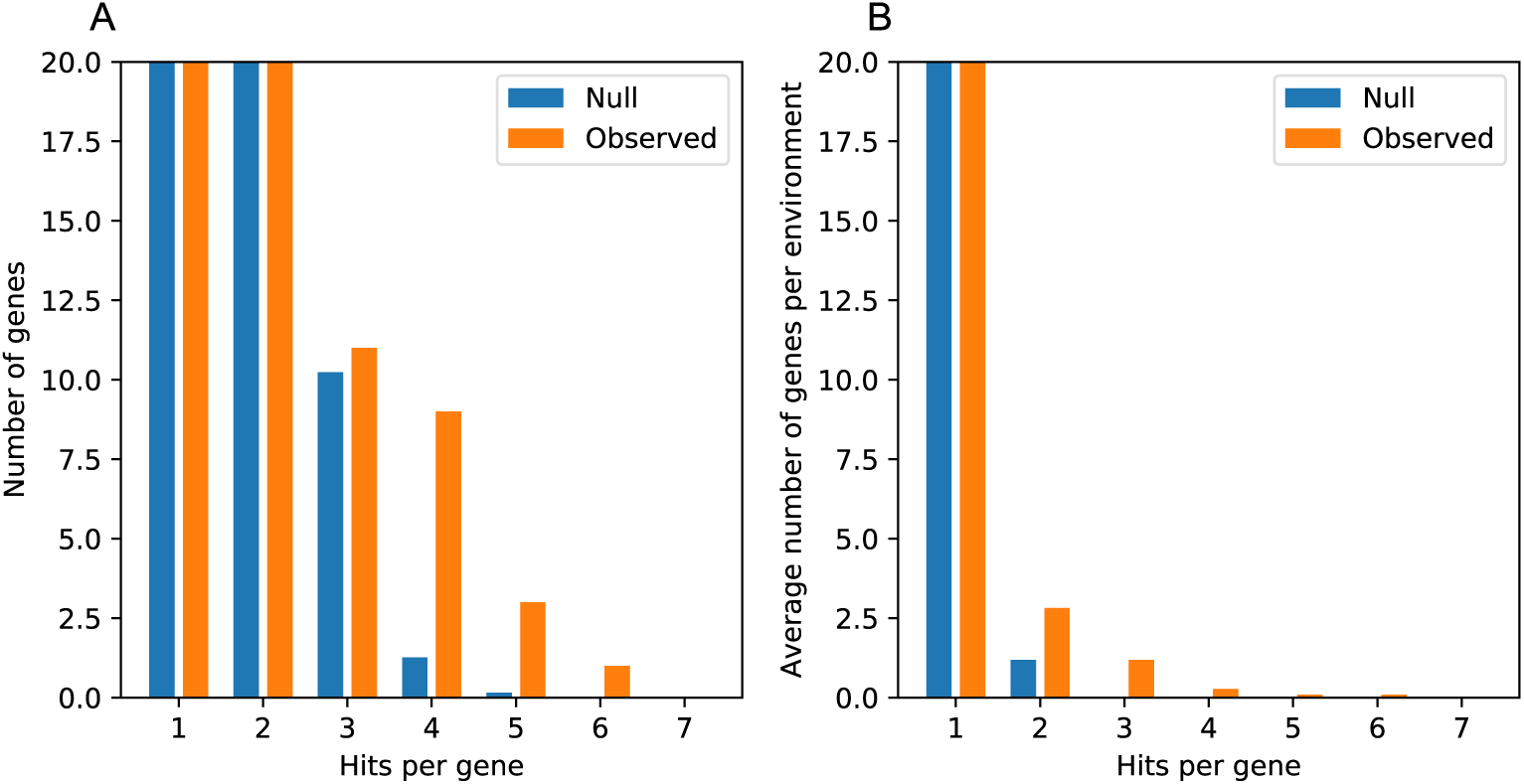
Enrichment of multi-hit genes across evolved lines. **A.** Number of genes with particular numbers of ‘hits,’ where a hit is defined as a nonsynonymous mutation in an independent clone. The null distribution represents all nonsynonymous mutations redistributed amongst the yeast genes, under a multinomial distribution with probability proportional to the gene length. Note that the bars showing genes with 1 or 2 hits are cut-off to more clearly show the differences in the rest of the distribution. **B.** Same as in A, where numbers of hits were counted within each environment, and the results were averaged across the environments. Note that the bars showing genes with one hit are cut-off to more clearly show the differences in the rest of the distribution.

**Fig S6.**
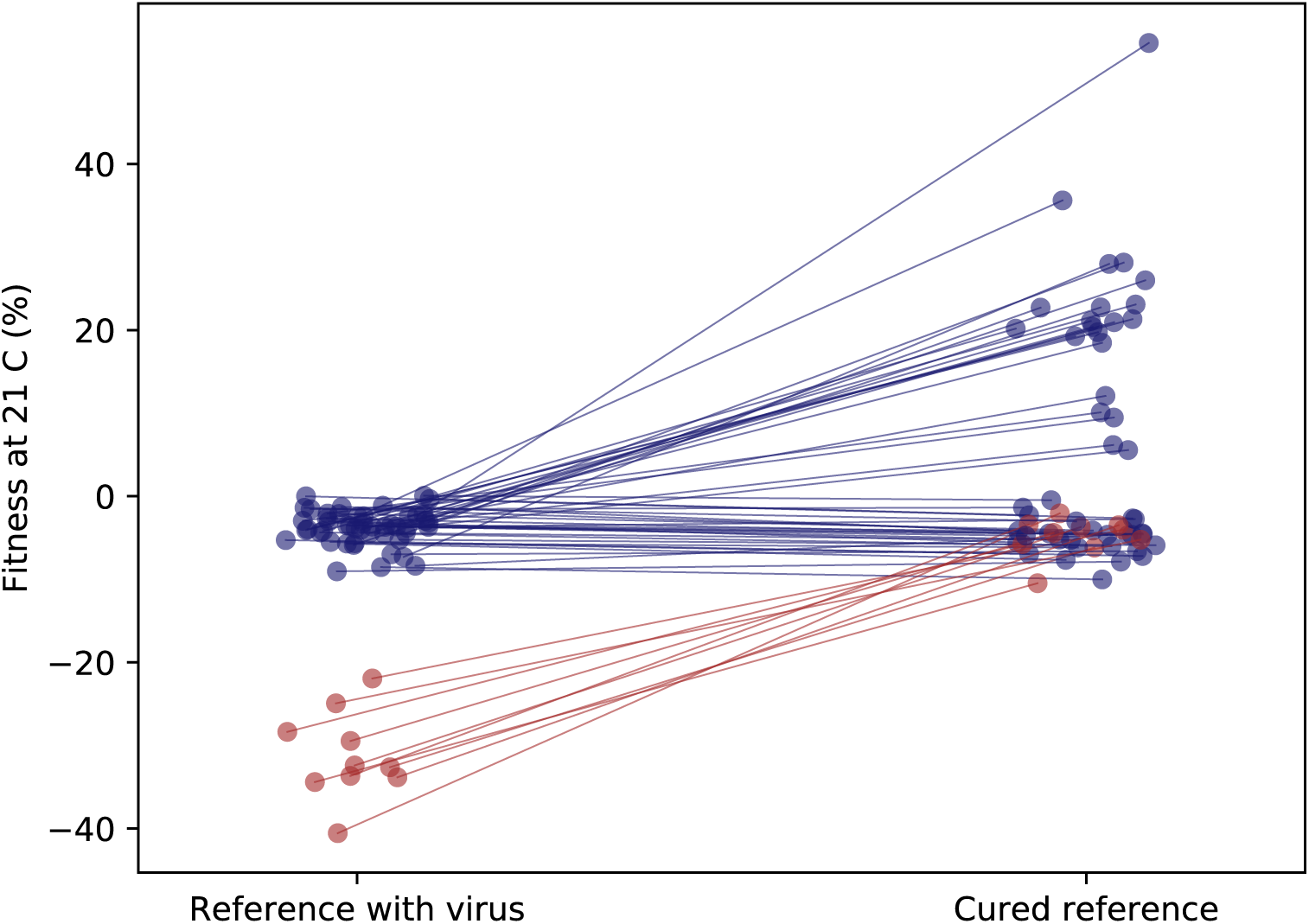
Fitness of clones relative to the original and virus-cured ancestor. As in Figure 3B, but for clones from environments Low Salt, pH 3.8, pH 6 that were not included in the diagnostic panel. Red clones were classified as V*^−^*, blue as V^+^.

**Fig S7.**
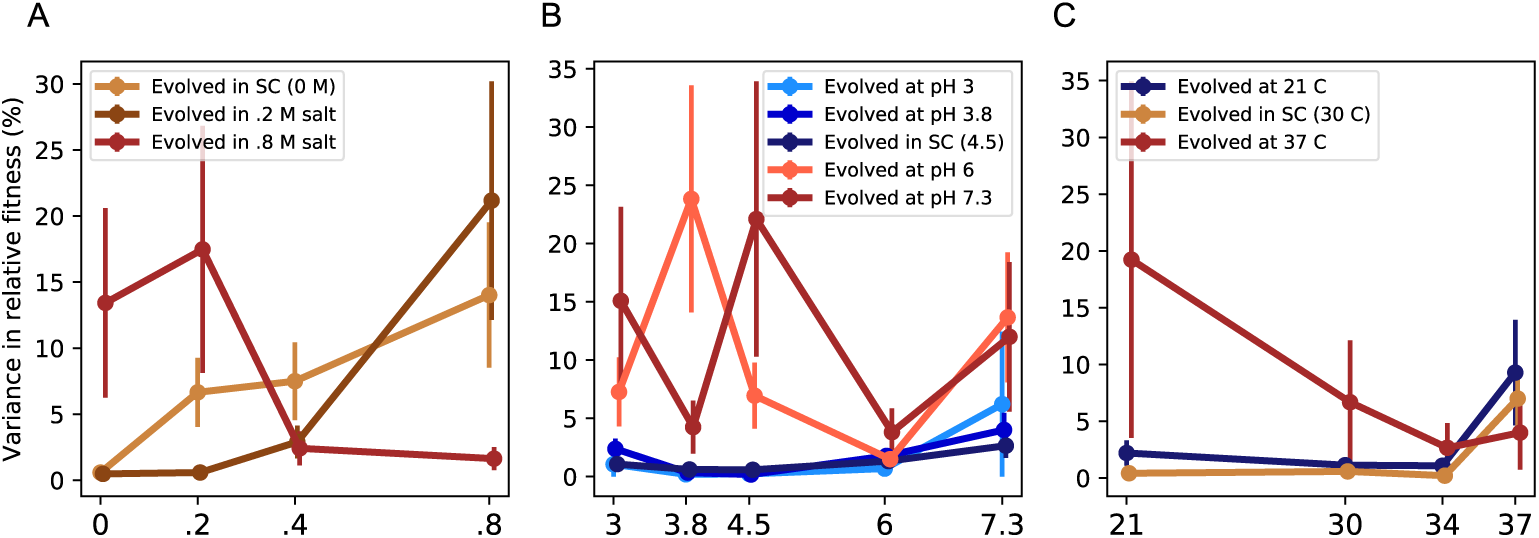
Variance in fitness across environmental panels. As in Figure 5A-C, but variance in fitness across groups of clones rather than means. Error bars represent ±1 standard error of the variance.

**Fig S8.**
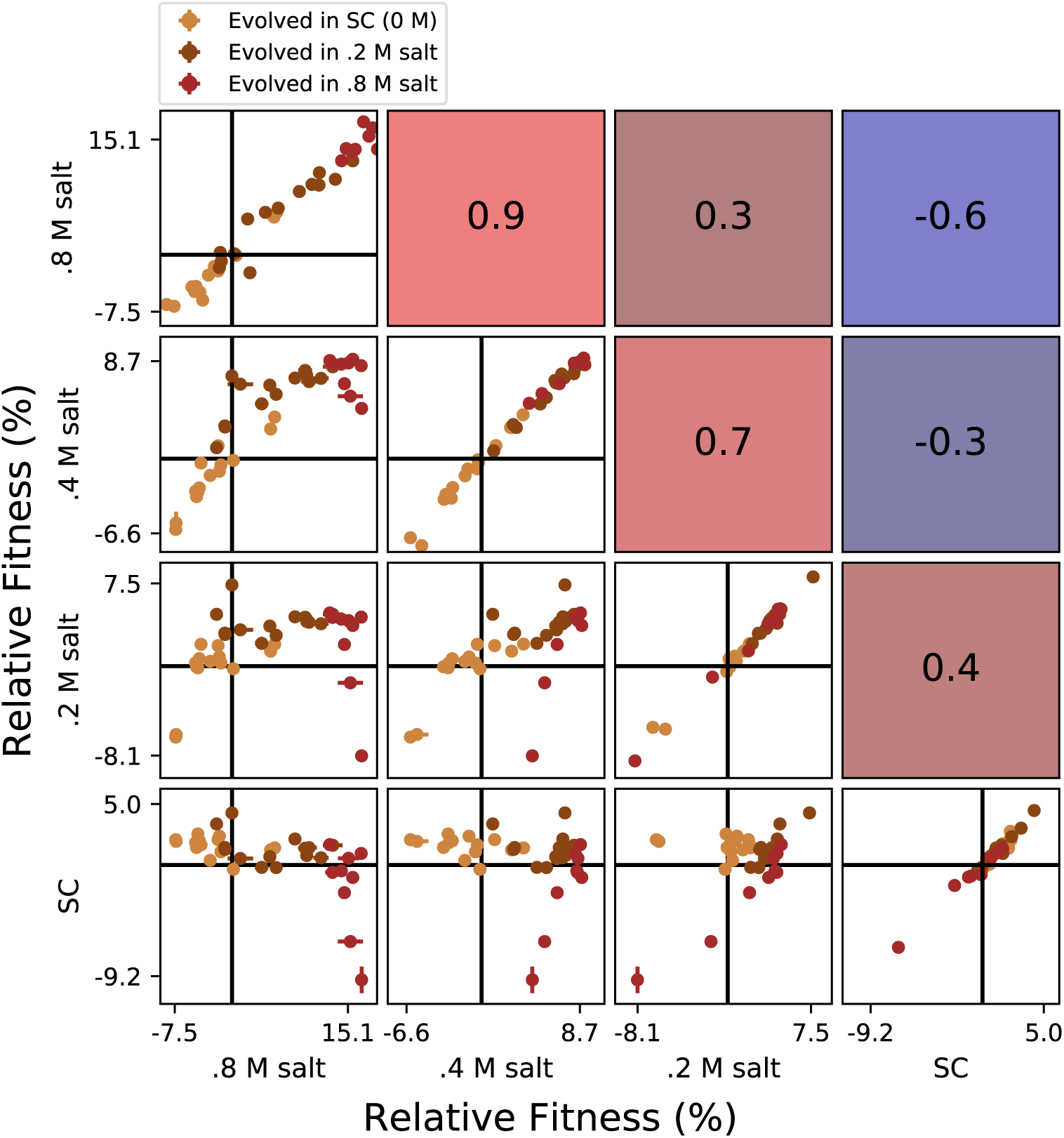
Correlations between clone fitness in different salt conditions. Each panel below the diagonal shows clone fitness in a particular pair of environments. (Error bars: ±1 SE on clone fitness.) The diagonal shows the correlation between technical replicates in the fitness assay in each condition. Panels above the diagonal are colored by and display the Pearson correlation coefficient between clone fitness in the corresponding pair of environments.

**Fig S9.**
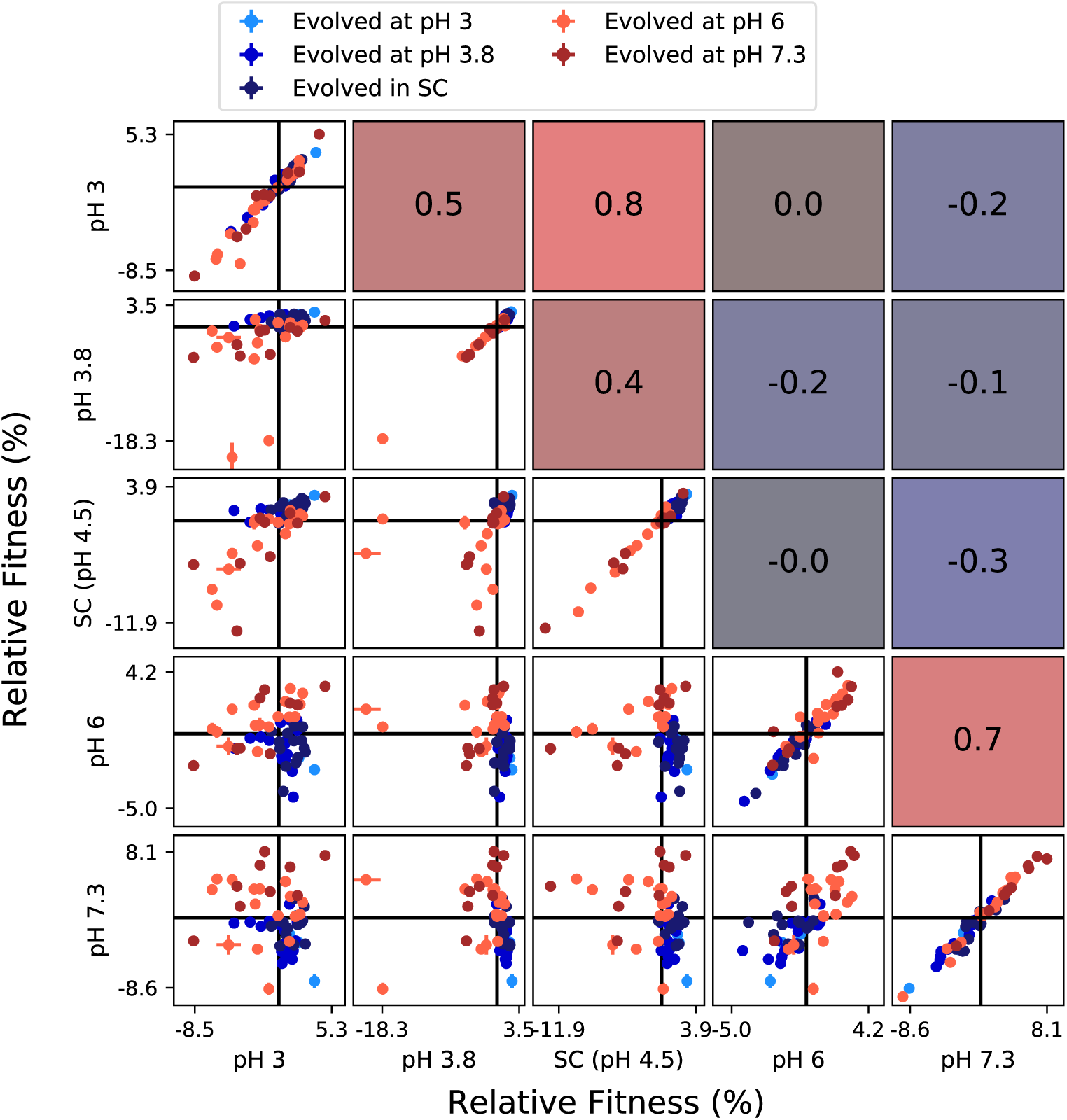
Correlations between clone fitness in different pH conditions. Each panel below the diagonal shows clone fitness in a particular pair of environments. (Error bars: ±1 SE on clone fitness.) The diagonal shows the correlation between technical replicates in the fitness assay in each condition. Panels above the diagonal are colored by and display the Pearson correlation coefficient between clone fitness in the corresponding pair of environments.

**Fig S10.**
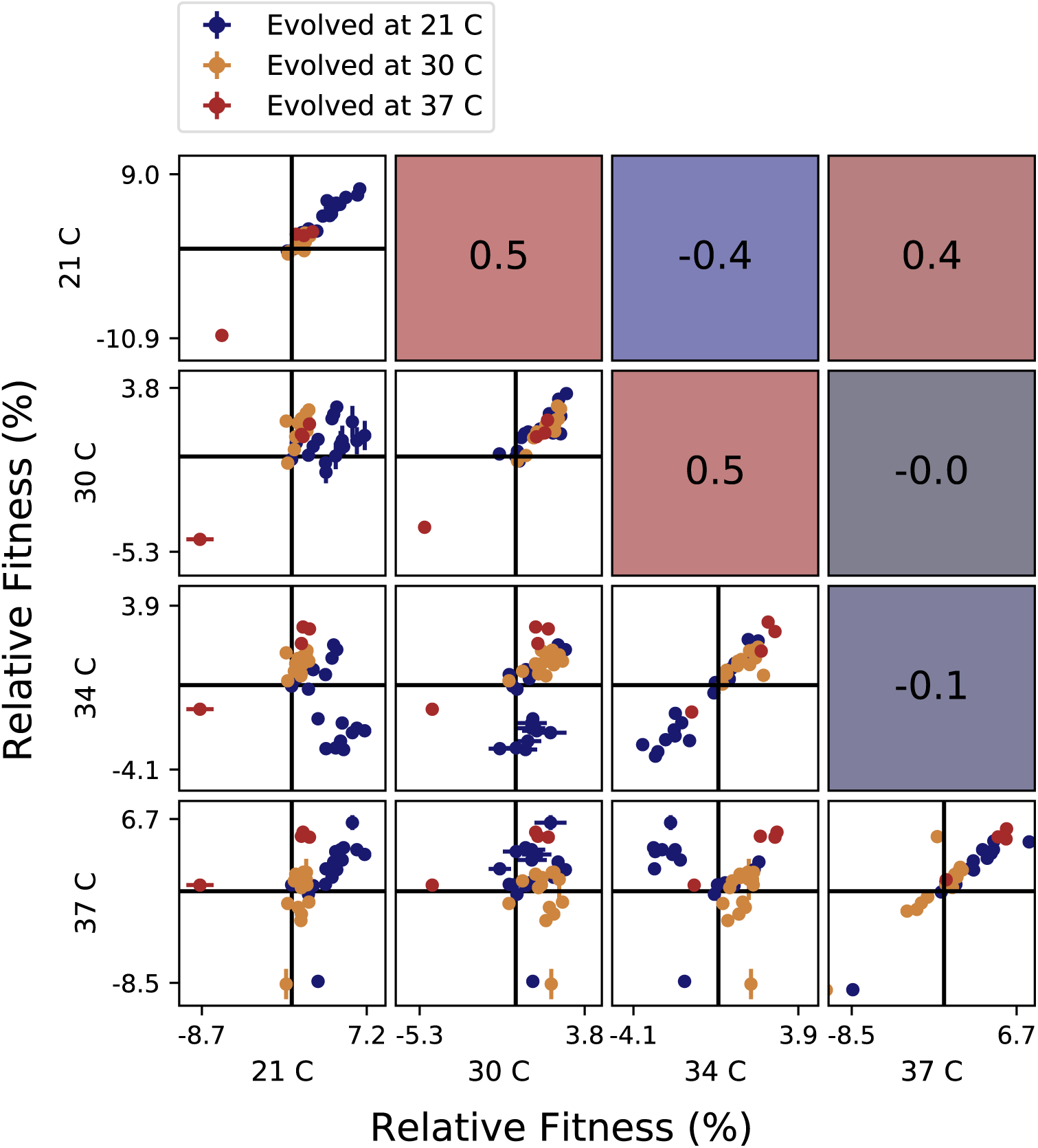
Correlations between clone fitness in different temperature conditions. Each panel below the diagonal shows clone fitness in a particular pair of environments. (Error bars: ±1 SE on clone fitness.) The diagonal shows the correlation between technical replicates in the fitness assay in each condition. Panels above the diagonal are colored by and display the Pearson correlation coefficient between clone fitness in the corresponding pair of environments.

**Fig S11.**
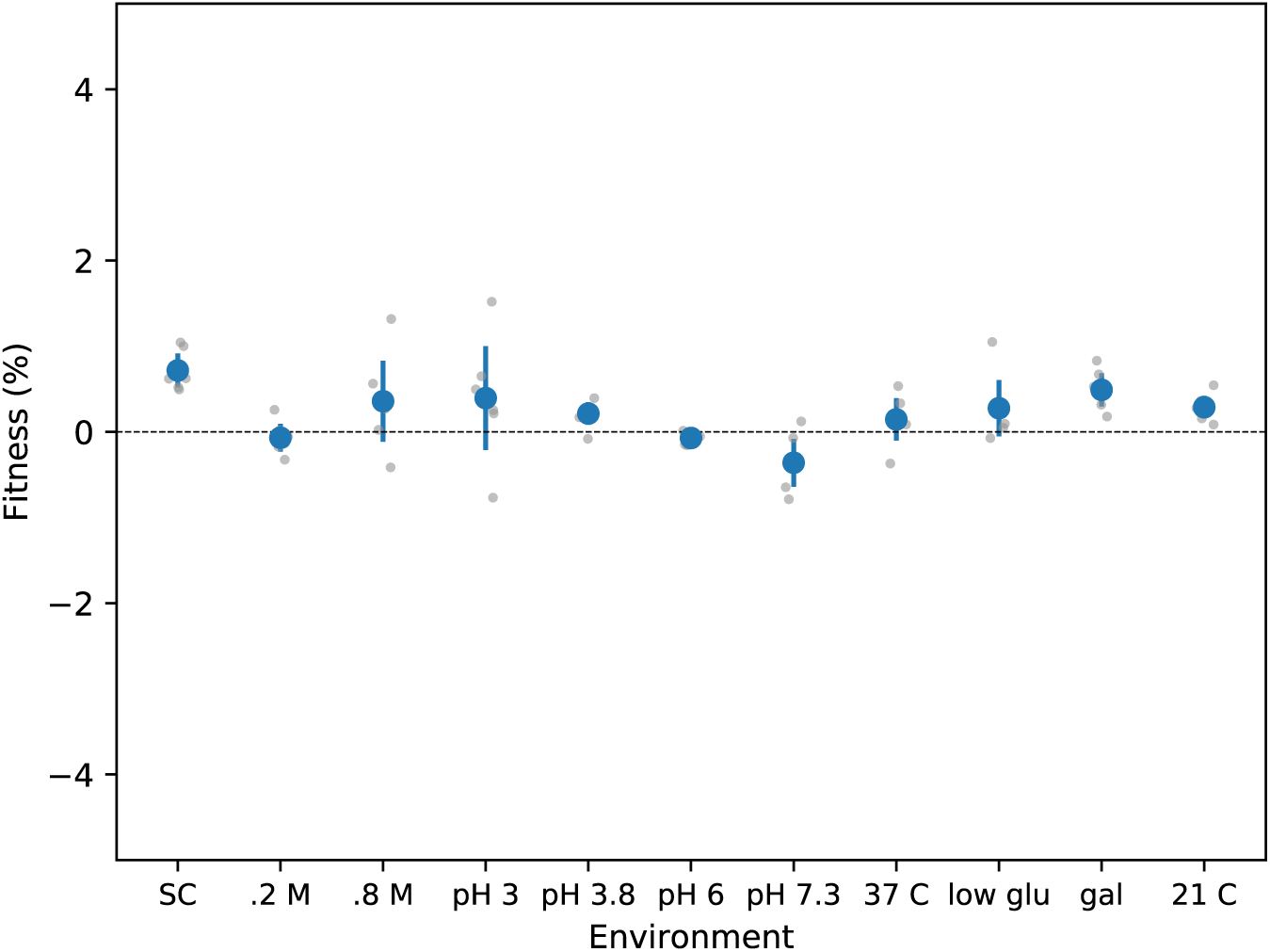
The fitness of the unlabeled ancestor, measured against the citrine-labeled ancestor. Fitness measurements were performed in 6 replicates (grey points); error bars represent *±* 2 SEM.

## References

1. Barrick JE, Lenski RE. Genome dynamics during experimental evolution. Nat Rev Genet. 2013;14(12):827–839.

2. Lee CE. Evolutionary genetics of invasive species. Trends Ecol Evol. 2002;17(8):386–391.

3. Bedhomme S, Hillung J, Elena SF. Emerging viruses: why they are not jacks of all trades? Curr Opin Virol. 2015;10:1–6.

4. Kassen R. The experimental evolution of specialists, generalists, and the maintenance of diversity. J Evol Biol. 2002;15(2):173–190.

5. Schluter D. Evidence for ecological speciation and its alternative. Science. 2009;323(5915):737–741.

6. Forister M, Dyer LA, Singer M, Stireman III JO, Lill J. Revisiting the evolution of ecological specialization, with emphasis on insect–plant interactions. Ecology. 2012;93(5):981–991.

7. Mitchell-Olds T, Willis JH, Goldstein DB. Which evolutionary processes influence natural genetic variation for phenotypic traits? Nat Rev Genet. 2007;8(11):845.

8. Savolainen O, Lascoux M, Merilä J. Ecological genomics of local adaptation. Nat Rev Genet. 2013;14(11):807–820.

9. Minor PD. Live attenuated vaccines: historical successes and current challenges. Virology. 2015;479:379–392.

10. Kim S, Lieberman TD, Kishony R. Alternating antibiotic treatments constrain evolutionary paths to multidrug resistance. Proc Natl Acad Sci USA. 2014;p. 14494–14499.

11. Imamovic L, Sommer MO. Use of collateral sensitivity networks to design drug cycling protocols that avoid resistance development. Sci Transl. 2013;5(204):204ra132.

12. Futuyma DJ, Moreno G. The evolution of ecological specialization. Annu Rev Ecol Syst. 1988;19(1):207–233.

13. Anderson JT, Willis JH, Mitchell-Olds T. Evolutionary genetics of plant adaptation. Trends Genet. 2011;27(7):258–266.

14. Bono LM, Smith Jr LB, Pfennig DW, Burch CL. The emergence of performance trade-offs during local adaptation: insights from experimental evolution. Mol Ecol. 2017;26(7):1720–1733.

15. Elena SF. Local adaptation of plant viruses: lessons from experimental evolution. Mol Ecol. 2017;26(7):1711–1719.

16. Levins R. Evolution in changing environments: some theoretical explorations. 2. Princeton University Press; 1968.

17. MacArthur RH. Geographical ecology: patterns in the distribution of species. Princeton University Press; 1984.

18. Stearns SC. Trade-offs in life-history evolution. Functional ecology. 1989;3(3):259–268.

19. Remold S. Understanding specialism when the jack of all trades can be the master of all. Proc Natl Acad Sci USA. 2012;279(1749):4861–4869.

20. Kawecki TJ. Accumulation of deleterious mutations and the evolutionary cost of being a generalist. Am Nat. 1994;144(5):833–838.

21. Whitlock MC. The red queen beats the jack-of-all-trades: the limitations on the evolution of phenotypic plasticity and niche breadth. Am Nat. 1996;148:S65–S77.

22. Bono LM, Draghi JA, Turner PE. Evolvability costs of niche expansion. Trends Genet. 2019;in press.

23. Anderson JT, Lee CR, Rushworth CA, Colautti RI, Mitchell-Olds T. Genetic trade-offs and conditional neutrality contribute to local adaptation. Mol Ecol. 2013;22(3):699–708.

24. Ågren J, Oakley CG, McKay JK, Lovell JT, Schemske DW. Genetic mapping of adaptation reveals fitness tradeoffs in *Arabidopsis thaliana*. Proc Natl Acad Sci USA. 2013;110(52):21077–21082.

25. Tiffin P, Ross-Ibarra J. Advances and limits of using population genetics to understand local adaptation. Trends Ecol Evol. 2014;29(12):673–680.

26. Cooper VS, Lenski RE. The population genetics of ecological specialization in evolving *Escherichia coli* populations. Nature. 2000;407(6805):736–739.

27. Turner PE, Elena SF. Cost of host radiation in an RNA virus. Genetics. 2000;156(4):1465–1470.

28. Zhong S, Khodursky A, Dykhuizen DE, Dean AM. Evolutionary genomics of ecological specialization. Proc Natl Acad Sci USA. 2004;101(32):11719–11724.

29. MacLean RC, Bell G, Rainey PB. The evolution of a pleiotropic fitness tradeoff in *Pseudomonas fluorescens*. Proc Natl Acad Sci USA. 2004;101:8072–8077.

30. Ostrowski EA, Rozen DE, Lenski RE. Pleiotropic effects of beneficial mutations in *Escherichia coli*. Evolution. 2005;59(11):2343–2352.

31. Duffy S, Turner PE, Burch CL. Pleiotropic costs of niche expansion in the RNA bacteriophage Φ6. Genetics. 2006;172:751–757.

32. Bennett AF, Lenski RE. An experimental test of evolutionary trade-offs during temperature adaptation. Proc Natl Acad Sci USA. 2007;104(Suppl 1):8649–8654.

33. Dettman JR, Sirjusingh C, Kohn LM, Anderson JB. Incipient speciation by divergent adaptation and antagonistic epistasis in yeast. Nature. 2007;447(7144):585.

34. Lee MC, Chou HH, Marx CJ. Asymmetric, bimodal trade-offs during adaptation of *Methylobacterium* to distinct growth substrates. Evolution. 2009;63(11):2816–2830.

35. Wenger JW, Piotrowski J, Nagarajan S, Chiotti K, Sherlock G, Rosenzweig F. Hunger artists: yeast adapted to carbon limitation show trade-offs under carbon sufficiency. PLoS Genet. 2011;7(8):e1002202.

36. Jasmin JN, Dillon MM, Zeyl C. The yield of experimental yeast populations declines during selection. Proc Natl Acad Sci USA. 2012;279(1746):4382–4388.

37. Jasmin JN, Zeyl C. Evolution of pleiotropic costs in experimental populations. J Evol Biol. 2013;26:1363–1369.

38. Yi X, Dean AM. Bounded population sizes, fluctuating selection and the tempo and mode of coexistence. Proc Natl Acad Sci USA. 2013;110(42):16945–16950.

39. Hietpas RT, Bank C, Jensen JD, Bolon DN. Shifting fitness landscapes in response to altered environments. Evolution. 2013;67(12):3512–3522.

40. Hong KK, Nielsen J. Adaptively evolved yeast mutants on galactose show trade-offs in carbon utilization on glucose. Metab eng. 2013;16:78–86.

41. Rodŕıguez-Verdugo A, Carrillo-Cisneros D, González-González A, Gaut BS, Bennett AF. Different tradeoffs result from alternate genetic adaptations to a common environment. Proc Natl Acad Sci USA. 2014;111(33):12121–12126.

42. Schick A, Bailey SF, Kassen R. Evolution of fitness trade-offs in locally adapted populations of *Pseudomonas fluorescens*. Am Nat. 2015;186(S1):S48–S59.

43. Leiby N, Marx CJ. Metabolic erosion primarily through mutation accumulation, and not tradeoffs, drives limited evolution of substrate specificity in *Escherichia coli*. PLoS Biol. 2014;12(2):e1001789.

44. McGee LW, Aitchison EW, Caudle SB, Morrison AJ, Zheng L, Yang W, et al. Payoffs, not tradeoffs, in the adaptation of a virus to ostensibly conflicting selective pressures. PLoS Genet. 2014 Oct;10(10):e1004611.

45. Fraebel DT, Mickalide H, Schnitkey D, Merritt J, Kuhlman TE, Kuehn S. Environment determines evolutionary trajectory in a constrained phenotypic space. eLife. 2017;6:e24669.

46. Lalić J, Cuevas JM, Elena SF. Effect of host species on the distribution of mutational fitness effects for an RNA virus. PLoS Genetics. 2011;7(11):e1002378.

47. Li C, Zhang J. Multi-environment fitness landscapes of a tRNA gene. Nat Ecol Evol. 2018;2(6):1025.

48. Selmecki AM, Maruvka YE, Richmond PA, Guillet M, Shoresh N, Sorenson AL, et al. Polyploidy can drive rapid adaptation in yeast. Nature. 2015;519(7543):349.

49. Gerstein AC, Chun HJE, Grant A, Otto SP. Genomic convergence toward diploidy in *Saccharomyces cerevisiae*. PLoS Genet. 2006;2(9):e145.

50. Venkataram S, Dunn B, Li Y, Agarwala A, Chang J, Ebel ER, et al. Development of a comprehensive genotype-to-fitness map of adaptation-driving mutations in yeast. Cell. 2016;166(6):1585–1596.

51. Voordeckers K, Kominek J, Das A, Espinosa-Cantú A, De Maeyer D, Arslan A, et al. Adaptation to high ethanol reveals complex evolutionary pathways. PLoS Genet. 2015;11(11):e1005635.

52. Harari Y, Ram Y, Kupiec M. Frequent ploidy changes in growing yeast cultures. Curr genet. 2018;64(5):1001–1004.

53. Wickner RB. Double-stranded and single-stranded RNA viruses of *Saccharomyces cerevisiae*. Annu Rev Microbiol. 1992;46(1):347–375.

54. Vagnoli P, Musmanno RA, Cresti S, Di Maggio T, Coratza G. Occurrence of killer yeasts in spontaneous wine fermentations from the Tuscany region of Italy. Appl Environ Microbiol. 1993;59(12):4037–4043.

55. Schmitt MJ, Breinig F. Yeast viral killer toxins: lethality and self-protection. Nat Rev Microbiol. 2006;4(3):212.

56. Greig D, Travisano M. Density-dependent effects on allelopathic interactions in yeast. Evolution. 2008;62(3):521–527.

57. Pieczynska MD, de Visser JAG, Korona R. Incidence of symbiotic dsRNA killer viruses in wild and domesticated yeast. FEMS yeast research. 2013;13(8):856–859.

58. Kandel JS. Killer systems and pathogenic yeasts. In: Koltin Y, Leibowitz MJ, editors. Viruses of fungi and simple eukaryotes. CRC Press; 1988. p. 243–263.

59. Schmitt MJ, Breinig F. The viral killer system in yeast: from molecular biology to application. FEMS Microbiol Reviews. 2002;26(3):257–276.

60. Buskirk SW, Rokes AB, Lang GI. Adaptive evolution of a rock-paper-scissors sequence along a direct line of descent. bioRxiv. 2019;p. 700302. https://www.biorxiv.org/content/10.1101/700302v1.

61. Liu H, Zhang J. Yeast Spontaneous Mutation Rate and Spectrum Vary with Environment. Curr Biol. 2019;29:1584–1591.

62. Sexton JP, Montiel J, Shay JE, Stephens MR, Slatyer RA. Evolution of ecological niche breadth. Annu Rev Ecol Evol Syst. 2017;48:183–206.

63. Good BH, Rouzine IM, Balick DJ, Hallatschek O, Desai MM. Distribution of fixed beneficial mutations and the rate of adaptation in asexual populations. Proc Natl Acad Sci USA. 2012;109(13):4950–4955.

64. Tikhonov M, Kachru S, Fisher DS. Modeling the interplay between plastic tradeoffs and evolution in changing environments. bioRxiv. 2019;https://www.biorxiv.org/content/10.1101/711531.

65. Lang GI, Murray AW. Estimating the per-base-pair mutation rate in the yeast *Saccharomyces cerevisiae*. Genetics. 2008;178(1):67–82.

66. Lang GI, Botstein D, Desai MM. Genetic variation and the fate of beneficial mutations in asexual populations. Genetics. 2011;188(3):647–661.

67. Kryazhimskiy S, Rice DP, Jerison ER, Desai MM. Global epistasis makes adaptation predictable despite sequence-level stochasticity. Science. 2014;344(6191):1519–1522.

68. Baym M, Kryazhimskiy S, Lieberman TD, Chung H, Desai MM, Kishony R. Inexpensive multiplexed library preparation for megabase-sized genomes. PLoS One. 2015;10(5):e0128036.

69. Jerison ER, Kryazhimskiy S, Mitchell JK, Bloom JS, Kruglyak L, Desai MM. Genetic variation in adaptability and pleiotropy in budding yeast. eLife. 2017;6:e27167.

70. Lang GI, Rice DP, Hickman MJ, Sodergren E, Weinstock GM, Botstein D, et al. Pervasive genetic hitchhiking and clonal interference in forty evolving yeast populations. Nature. 2013;500(7464):571–574.

71. Sherman F, Fink GR, Hicks JB. Methods in yeast genetics: laboratory manual. Cold Spring Harbor, New York: Cold Spring Harbor Laboratory; 1981.

72. Storici F, Lewis LK, Resnick MA. In vivo site-directed mutagenesis using oligonucleotides. Nat Biotech. 2001;19(8):773–776.

